# Essentiality and dynamic expression of the human tRNA pool during viral infection

**DOI:** 10.1101/2024.04.25.591047

**Authors:** Noa Aharon-Hefetz, Michal Schwartz, Orna Dahan, Noam Stern-Ginossar, Yitzhak Pilpel

**Affiliations:** Department of Molecular Genetics, Weizmann Institute of Science, Rehovot, 76100, Israel

## Abstract

Human viruses depend on the translation resources of the host cell. A significant translation resource is the tRNA pool of the cell, as human viruses do not encode tRNA genes. Through tRNA sequencing, we inspected the human tRNA pool upon infection of human Cytomegalovirus (HCMV) and SARS-CoV-2. HCMV-induced alterations in tRNA expression were predominantly virus-driven, with minimal influence from the cellular immune response. Notably, specific tRNA post-transcriptional modifications appeared to modulate stability and were susceptible to HCMV manipulation. In contrast, SARS-CoV-2 infection did not significantly impact tRNA expression or modifications. We compared the codon usage of viral genes to the proliferation-differentiation codon-usage signatures of human genes. We found a marked difference between the viruses, with HCMV genes aligning with differentiation codon usage and SARS-CoV-2 genes reflecting proliferation codon usage. We further found that codon usage of structural and gene expression-related viral genes displayed high adaptation to host cell tRNA pools. Through a systematic CRISPR screen targeting human tRNA genes and modification enzymes, we identified specific tRNAs and enzymes that improve or reduce HCMV infectivity and cellular growth. These findings highlight the dynamic interplay between the tRNA pool and viral infection dynamics, shedding light on mechanisms governing host-virus interactions.

## Introduction

A significant challenge for viruses upon cellular infection is to execute their gene expression programs. Viruses must efficiently and promptly translate a substantial portion of their proteins to facilitate effective infection while coping with the host cell’s anti-viral response. Unlike bacterial viruses (Guerrero-Bustamante & Hatfull, 2024; Limor-Waisberg et al., 2011), human viruses do not encode tRNA genes (Albers & Czech, 2016). Consequently, human viruses rely on the host’s tRNA pool and associated proteins to fulfill the translation needs for virus propagation, necessitating the hijacking of the host’s translation machinery (Hoang et al., 2021; Rozman et al., 2023).

Cells evolved to coordinate the supply of the tRNA pool to the codon demand of the transcriptome (Boël et al., 2016; Hanson & Coller, 2018), allowing cells to achieve efficient and accurate translation (Frumkin et al., 2018; Konrad L M Rudolph et al., 2016). Viruses employ various strategies to influence the translation system in favor of expressing their own genes, including adapting to the codon usage of their host genes (Bahir et al., 2009; Hernandez-Alias et al., 2021) or disrupting the natural equilibrium between tRNA availability and codon demand (Pavon-Eternod, David et al., 2013; Van Weringh et al., 2011).

The manipulation of the tRNA pool can occur through the regulation of tRNA gene transcription and alterations in tRNA modification levels (Pan, 2018). Among cellular RNAs, tRNAs exhibit extensive post-transcriptional modifications, displaying specificity to the type of modification and nucleotide positions within the tRNA molecules (Lucas et al., 2024; Suzuki, 2021). Modifications of tRNAs can affect their RNA stability (De Zoysa et al., 2024; Wang et al., 2023) and decoding (Giguère et al., 2024; Krueger et al., 2024; Saleh & Farabaugh, 2024), among others. Previous research has provided experimental evidence highlighting the significance of tRNA modification in regulating viral infection. For instance, experiments involving the knock-out of TYW1, one of the Wybutosine modification ‘writers’, demonstrated an increased ribosomal frameshift in a reporter construct containing the frameshift site of the HIV gag-pol slippery codon motif. This elevation in ribosomal frameshift could impact the crucial gag-pol ratio, influencing the loading of reverse transcriptase into new viral capsids and thereby affecting viral progeny (Rak et al., 2021; Rosselló-Tortella et al., 2020)

In recent years, we have seen significant advancement in measuring and quantifying changes in amounts and modification levels of the tRNA pool in diverse processes (Behrens et al., 2021; Zheng et al., 2015). These means can potentially improve the resolution in which we capture dynamic changes in the tRNA pool of viral infected cells. In human cells, tRNA sequencing enabled the detection of a significant transition in the tRNA pool upon switch to normal or malignant transformation proliferation (Gingold et al., Rak et al., 2021; Santos et al., 2019; Zhang et al., 2018) between different tissues (Z. Zhang et al., 2018) and in various stress conditions (Ling et al., 2014; Mahalingam et al., 2022). Furthermore, tRNA sequencing allows accurate monitoring of changes in tRNA modification levels (W. Zhang et al., 2022).

Yet, passive monitoring of dynamics in the tRNA pool, without manipulating it, falls short of deducing functional effects. Indeed, a first step was introduced towards a systematic knock-out of 20 human tRNA families in a pooled fashion using CRISPR-Cas9 (Aharon-Hefetz et al., 2020). That screen revealed the essential role of specific tRNA genes for cell proliferation and others for cellular arrest. tRNA essentiality showed limited agreement with tRNA expression changes during such conditions (Aharon-Hefetz et al., 2020).

In the present study, we utilized tRNA sequencing to observe alterations in the quantity and epi-transcriptomic modifications of all human tRNA genes following infection with HCMV and SARS-CoV-2. We identified changes in the expression and modification levels of tRNA genes in HCMV-infected cells, while the tRNA pool of SARS-CoV-2 infected cells remained stable following infection. Treating uninfected cells with interferon (IFN), we revealed a mild effect of the anti-viral response on the tRNA pool. A comparison between the codon usage of viral genes and the proliferation-differentiation codon usage of human genes revealed a differential adaptation of HCMV and SARS-CoV-2 genes to the signatures of differentiation and proliferation codon usage, respectively. However, structural and gene expression-related genes of both viruses showed the highest adaptation to the tRNA pool of infected cells. To systematically investigate the functional essentiality of all human tRNAs for HCMV infection, encompassing cytosolic tRNA genes, pseudo tRNA genes, and tRNA modification enzymes, we generated a comprehensive library of 3000 sgRNAs for CRISPR screening. This library allowed us to conduct knock-out screenings under normal cellular growth conditions and during HCMV infection. It pinpointed specific tRNAs and modification enzymes needed or restricted to either of these conditions. This study represents one of the initial explorations into elucidating the fundamental role of tRNAs in both cellular viability and viral infection.

## Results

### tRNA levels are modulated in response to HCMV infection

We aimed to explore tRNAs’ dynamic change during viral infections. We first examined the tRNA pool of HCMV-infected human foreskin fibroblasts (HFFs) cells. HCMV has a prolonged replication cycle, with genome replication starting around 20 hours post-infection and beginning progeny release at around 4 days post-infection. We thus sampled the tRNA pool at 5, 16, 24, and 72 hours post-infection (hpi). Comparing the tRNA pool of uninfected cells to infected cells, we observed marked differences in the expression of many cytosolic tRNA isodecoder genes starting from 24hpi and becoming more pronounced at 72hpi (Figure 1A-C). Specifically, most of the lowly expressed tRNAs in uninfected cells are up-regulated during infection (Figure 1C).

**Figure 1.**
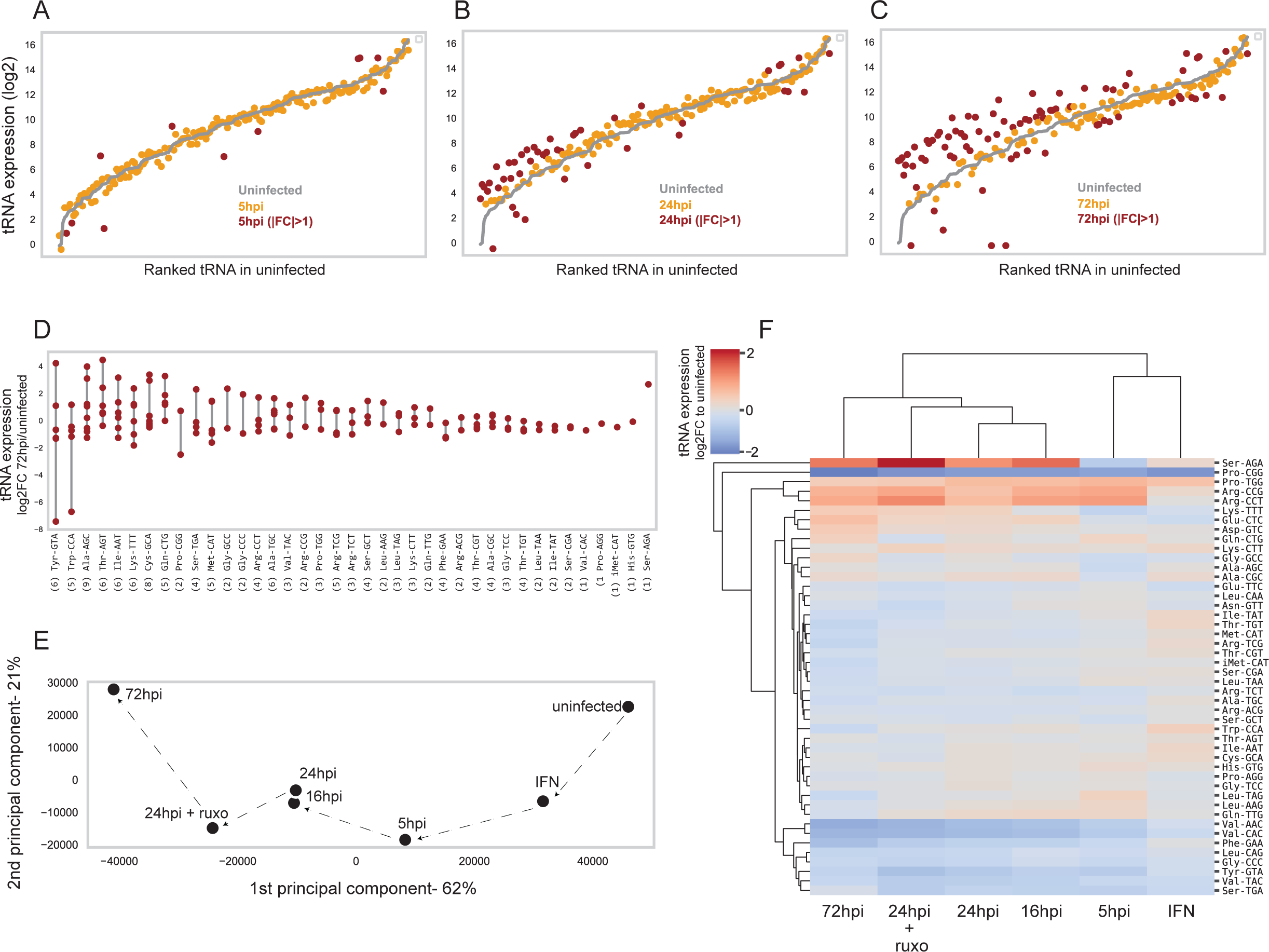
-changes in the tRNA expression of HCMV-infected HFFs A-C| Cytosolic tRNA expression (log2) of uninfected HFFs (in gray lines) and HCMV-infected cells (in yellow-red) in a time-course manner. The tRNAs are ordered based on their expression level in uninfected cells. tRNA genes that differentially expressed (|FC|>1) are marked in dark red. A| 5 hours post-infection (hpi); B| 24 hpi; C|72 hpi. The tRNA pool of 16 hpi is highly similar to that of 24 hpi, thus not shown here. D| Differences in the expression level of tRNA isodecoders. The log2 fold-change in expression at 72 hpi relative to uninfected cells of each tRNA isodecoder is presented for each tRNA isoacceptor. The numbers in parenthesis depict the number of isodecoders presented in each tRNA isoacceptor. E| Principle component analysis (PCA) of the HCMV-infected samples based on the mean tRNA expression of three biological repeats. The dashed arrows emphasize the gradual progress in infection. F| Hierarchical clustering of the HCMV-infected samples and tRNA families based on the averaged changes (two or three biological repeats) in the summed read fraction of each tRNA family relative to the uninfected sample (log2).

We further investigated the expression level of mitochondrial tRNAs. Given that the expression level of mitochondrial tRNAs is a cumulative sum of contributions from many mitochondria within each cell, we normalize it by the estimated mitochondrial genome copy number per cell, which is about 300 particles in human skin fibroblasts (Jiang et al., 2020). Per mitochondrial genome copies, the expression of mitochondrial tRNA is comparable to the lowest among the cytosolic tRNAs (Figure S1A). Furthermore, we observed a slight upregulation of the mitochondrial tRNA expression at 72hpi. Considering that host mitochondria biogenesis is increased following HCMV acute infection (Combs et al., 2020), the observed elevation in mitochondrial tRNAs could be due to increased mitochondrial units in the infected cells and changes in expression of specific genes. In most genomes, including humans, most tRNA families consist of several genes with the same anticodon, called tRNA isodecoders. It was previously shown that the expression of tRNA isodecoders might be differentially regulated between cell types and conditions (Gao et al., 2024; Sagi et al., 2016). As some tRNA isodecoders have unique sequences, we can detect potential differences in expression changes among such families through sequencing. Focusing on the cytosolic tRNAs, we observe a broad range of expression changes among tRNA isodecoders in response to HCMV infection (Figure 1D). In fact, the range spanned by some of the tRNA isodecoder families can be as broad as the change between families with different anticodons (Figure 1D).

The changes observed in the tRNA pool can be driven by the virus or might be governed by the host in response to infection. To explore a potential host response component in the absence of infection, we treated the cells with a mixture of IFNa+b, a treatment that mimics the response of cells to infection, yet without the virus. To further distinguish between the effect of the virus and the host response, we treated HCMV-infected cells with ruxolitinib, which blocks the native IFN response. Calculating the Pearson correlation between the tRNA pools upon repeats of the different time points following infection and treatments showed high consistency (Figure S1B, R >= 0.85). We used a clustergram and principal component analysis (PCA) based on averaged tRNA expression between repeats to present the similarity between the tRNA pools of cells at different time points along HCMV infection and under the abovementioned treatments. We observed gradual changes in the tRNA pool with the progression of the infection (Figure 1E). The most deviant tRNA pool was obtained at the end of the time course (72 hpi), in which the highest number of tRNAs showed a modified expression level relative to uninfected cells. The IFNa+b treated cells that were not infected showed a relatively small change in the tRNA pool compared to the control sample (i.e., untreated and uninfected cells) and in the same direction as the change following infection (Figure 1E). This indicates that the interferon response has a minor contribution to the shift in tRNA expression, which is thus deduced to be mediated mainly by the virus. The infected cells treated with ruxolitinib and then harvested at 24hpi are localized in expression space between 24hpi and 72hpi, indicating an improved infection in these cells. This result strengthens the indication that the virus is the entity that drives the change in tRNA expression and not the host cells in response to infection.

The clustegram shown in Figure 1F illustrates the similarity between samples based on the fold change in the expression of cytosolic tRNA isoacceptors during HCMV infection and the abovementioned treatments relative to control sample. Consistent with the PCA picture, the early infection sample (5hpi) is clustered with the IFN-treated sample, while the later time points are clustered separately. In 5hpi, a few tRNAs already changed their expression level, which intensified in later time points, e.g., Val-CAC tRNA. Most tRNAs do not respond, showing only mild changes in tRNA expression level, e.g., Glu-TTC tRNA. However, several tRNAs show a significant difference in expression. The most changing ones show up or down-regulation by a factor of up to 4, e.g., Arg-CCT and Val-AAC tRNAs, even within a day (Figure 1F). Human tRNAs are very stable, with a half-life of 100 hours (Choe & Taylor, 1972) in a normal decay process. Thus, even a putative transcription arrest cannot give rise to a 4-fold decline within 24 hours. Further, the cell cycle of infected cells is arrested (Bogdanow et al., 2021). Thus, dilution of the tRNAs through cell division cannot explain the reduction in tRNAs. Therefore, active degradation of these tRNAs during viral infection is the plausible means that could lead to such drastic down-regulation.

Specific tRNAs stand out in their dynamics. Ser-AGA is down-regulated in IFN and early infection, and it is then up-regulated more than four fold at later time points of infection. Pro-TGG shows the opposite behavior -its expression increases twofold in IFN and 5hpi samples, then decreases to basal levels in late infection. Pro-CGG expression is falling 4-fold in all samples. Interestingly, we found that their transcription level is stable during HCMV infection relative to uninfected cells, as was determined by deep-sequencing of their nascent transcripts (Mahalingam et al., 2022) (Figure S1C). These results suggest that post-transcription regulation induces expression level changes of several tRNAs in opposing directions, which could be driven by the virus or the host response.

### tRNA modifications are modulated in response to HCMV infection

We further examined and quantified the change in tRNA modifications during HCMV infection based on tRNA sequencing and the effects of modifications on the obtained sequence, as described by Rak et al., 2021. We found that the level of most of the modifications does not change between uninfected and late-infected samples (Figure 2A, r=0.95, p<0.001). Yet, we did find several modification types that show a significant change in specific tRNAs. Modifications in Val-CAC, Val-TAC, and Gly-CCC tRNA genes (dihydrouridine modifications that occur at positions 20, 20, and 19, respectively) were reduced in infected cells (Figure 2A). It was previously shown that dihydrouridine modification in tRNA genes contributes to the stability of the tRNA molecule (Faivre et al., 2021). Indeed, we observed a reduction in the levels of these tRNAs in late infection stages (Figure 1F). Thus, the reduced expression of Val-CAC, Val-TAC, and Gly-CCC tRNAs in late infection might be explained by the reduction in dihydrouridine modification via the effect on tRNA stability. However, we did not detect changes in dihydouridine modification on other tRNAs that harbor this modification in human cells, nor did we observe changes in their expression level. It is worth mentioning that Val-CAC, Val-TAC, and Gly-CCC tRNAs are the most highly expressed in uninfected cells among the tRNAs that have dihydrouridine modification (data not shown). These findings suggest that during HCMV infection, there is no global effect on the writers of dihydouridine modification, but rather, the changes in dihydrouridine modification levels are specific to highly-expressed tRNAs.

**Figure 2.**
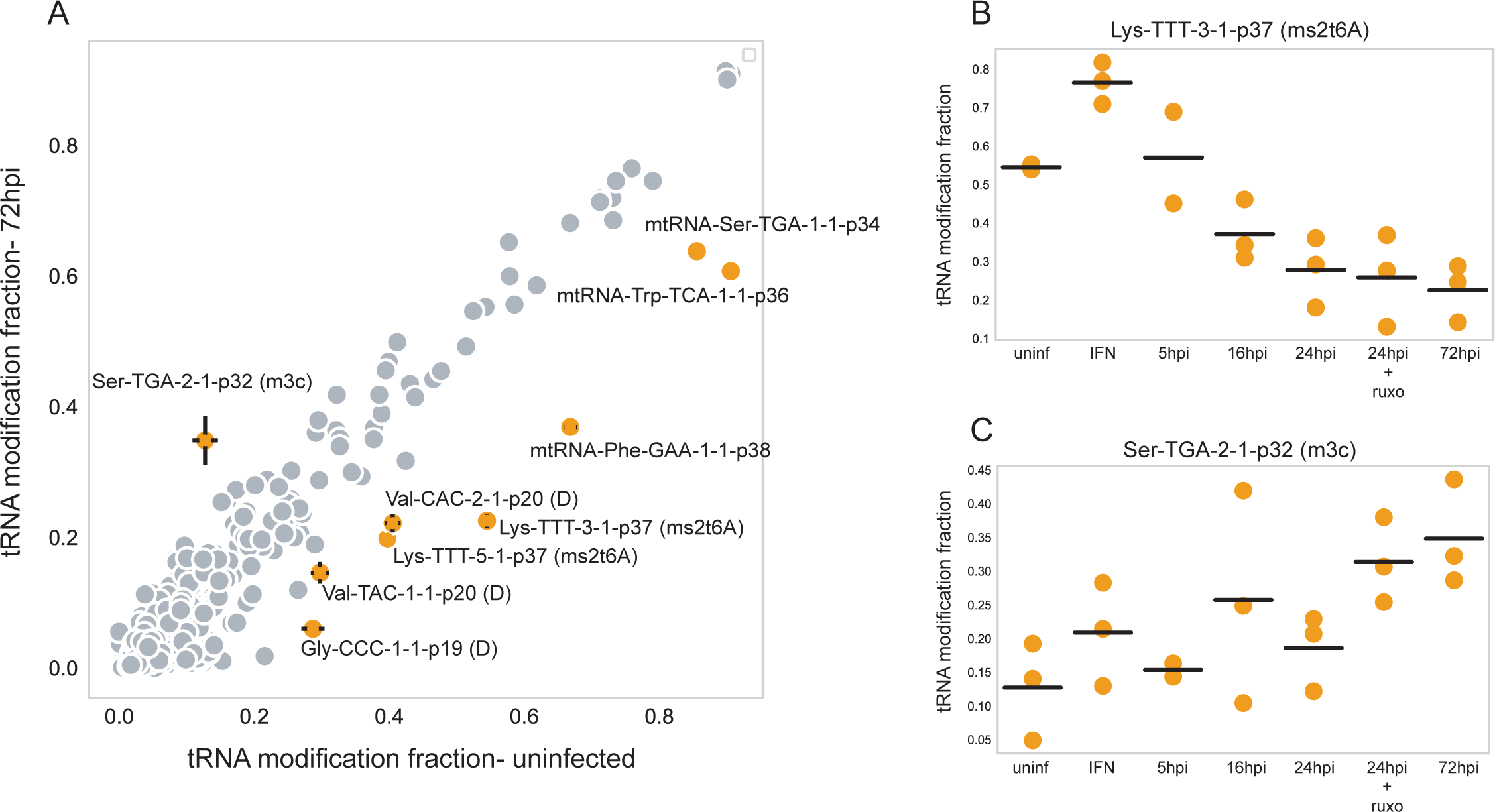
-changes in the tRNA modification levels of HCMV-infected HFFs A| A scatter plot describes the change in the averaged tRNA modification level in the uninfected sample (x-axis) relative to 72 hpi (y-axis). Modification levels significantly regulated following infection are marked in yellow with error bars showing the confidence interval calculated from three biological repeats and their name, including the names of the tRNA, the position in the tRNA gene, and the type of base modification. Pearson r = 0.95, p-value<0.001. B-C| Change in the tRNA modification level along HCMV infection on B| Lys-TTT-3-1 gene, position 37, modification ms2t6A; C| Ser-TGA-2-1, position 32, modification m3C. For each sample, each dot depicts a biological repeat (3 repeats in total) and the lines represent the averaged modification level.

We further explored the dynamic change in the level of modifications in the anticodon area that was shown to affect decoding in the ribosome (Rapino et al., 2017). Interestingly, the levels of ms2t6A on Lys-TTT genes at position 37 increased in IFN-treated cells, whereas along infection, it gradually decreased (Figure 2B and S2). On the other hand, the levels of the m3C modification on Ser-TGA-2-1 showed a mild up-regulation in IFN-treated cells relative to uninfected cells and a more substantial upregulation during HCMV infection (Figure 2C). We also found lower modification levels in the anticodon loop of mitochondrial tRNAs-Ser-TGA, Trp-TCA, and Phe-GAA (Figure 2A). Interestingly, these modifications were previously shown to be associated with cell proliferation (Ignatova et al., 2020; Rak et al., 2021; Rosselló-Tortella et al., 2020; Smith et al., 1985), suggesting a relationship with the cell cycling phase crucial for efficient HCMV infection (Bogdanow et al., 2021; Fortunato et al., 2002).

Altogether, these results suggest that the virus is the main contributor to the shifts in the tRNA pool. The anti-viral stress response has a minor impact, often aligning with that initiated by the virus. However, there are instances where the effects are opposite, affecting specific tRNA expression and modifications near the anticodon.

### Codon usage adaption of HCMV genes to the tRNA pool of infected cells

To examine the function of translation supply and demand between tRNAs and the codons in human-HCMV interaction, we computed and compared the codon usage of HCMV genes to that of human genes. A significant distinction in codon usage of mammalian genes is between the proliferation and cell-autonomous related functions and the differentiation and multicellularity programs (Gingold et al., 2014; Zviran et al., 2018; Hernandez-Alias et al., 2020). We scored each human and viral gene according to the resemblance of their codon usage to the codon usage signature of proliferation and differentiation-related genes in humans (Figure 3A-B and S3A). Focusing first on all human transcripts, we observe that 30% of the transcripts tend to have a higher correlation with the differentiation and multicellularity codon usage signatures (r > 0.8) than with the proliferation and cell-autonomous functionalities (r < 0.6). A smaller percentage of the transcripts (18%) have the opposite tendency, i.e., high codon usage similarity to the proliferation signature (r > 0.75) and low similarity to the differentiation signatures (r < 0.35). About 5% of human transcripts are lowly correlated with either of the two programs (differentiation: r < 0.6; proliferation: r < 0.6) (Figure S3A).

**Figure 3.**
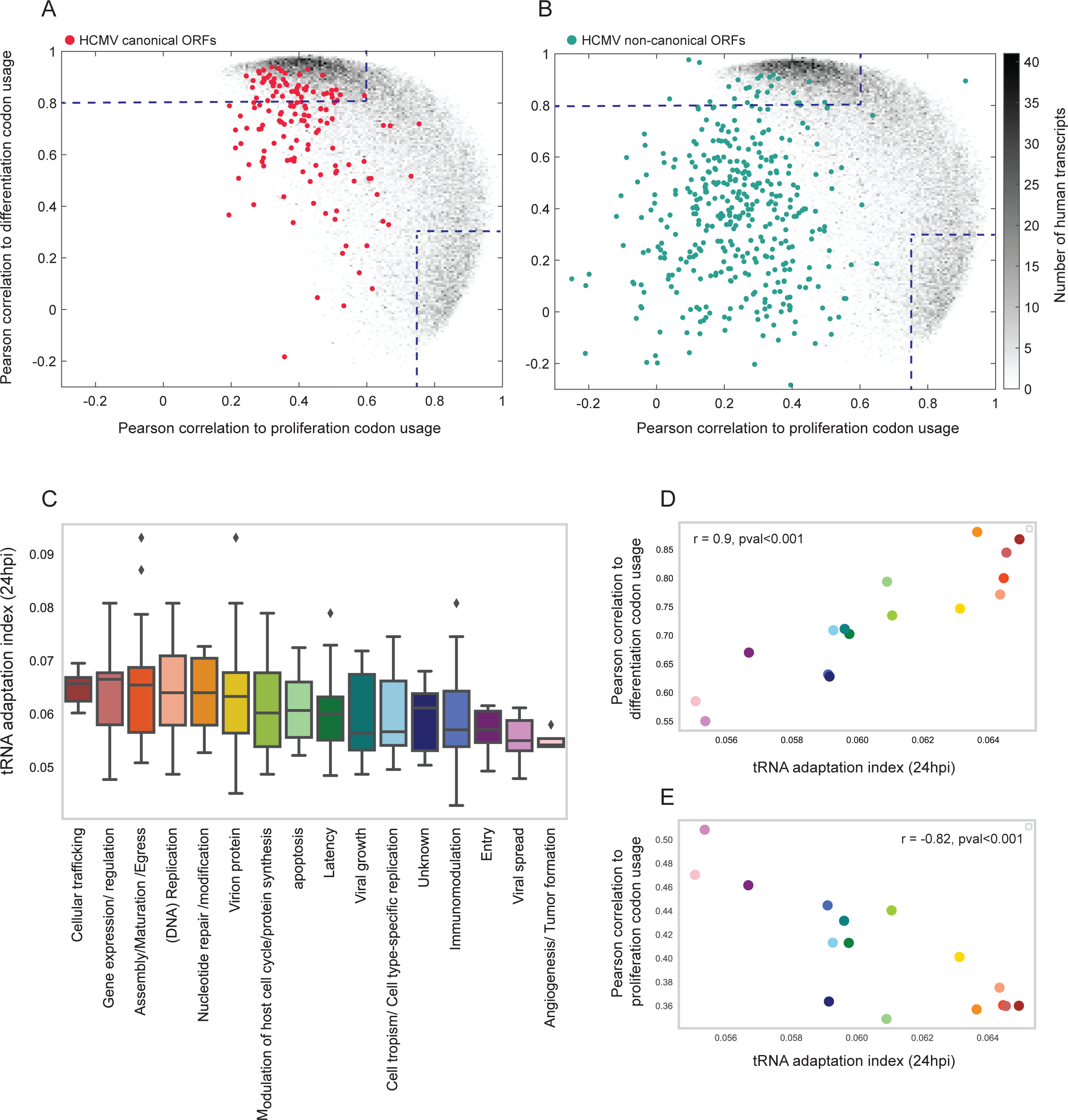
-Codon usage adaptation of HCMV ORFs to the tRNA pool of HCMV-infected HFFs A-B| Density plots of the Pearson correlation measured for each transcript to the human proliferation and differentiation codon-usage signatures. Gray dots denote human transcripts, whereas color dots denote HCMV ORFs. The color bar (gray) represents the number of human transcripts (36762). The dashed lines represent the Pearson coefficients that determine high similarity to the proliferation or differentiation codon-usage signatures. A| HCMV canonical ORFs (146), B| HCMV non-canonical ORFs (110). The analysis did not include short non-canonical ORFs (codon count < 58). C| A box plot depicting the tRNA adaptation index (tAI) of canonical HCMV ORFs as calculated based on the tRNA pool of 24hpi. HCMV ORFs are grouped according to their functionality, as was determined by Ye et al., 2020. D-E| Scatter plots showing the averaged tAI of functional groups of HCMV ORFs at 24hpi (x-axis) and their averaged Pearson correlation to D| differentiation codon usage (Pearson r = 0.9, p-value < 0.001), E| proliferation codon usage (Pearson r = -0.82, p-value < 0.001) (y-axis). The color code of each functional category corresponds to Figure 3C.

Inspecting the HCMV “canonical” open reading frames (ORFs), i.e., the 146 ORFs whose expression is regulated in a temporal cascade following infection (Rozman et al., 2022; Stern-Ginossar et al., 2012), we found that 39% of them have a codon usage signature that correlates more with the differentiation signature (r > 0.8) than with the proliferation signature (r < 0.6). Interestingly, none of the HCMV canonical ORFs are highly correlated with the proliferation signature (r > 0.75) (Figure 3A).

In HCMV-infected cells, viral gene expression does not initiate if the cells are actively replicating (Fortunato et al., 2002), and the virus is known to modulate the cell cycle to establish a unique cell cycle arrest at the G1/S transition (Bogdanow et al., 2021). Thus, the codon usage adaptation of HCMV to the differentiation signature and away from the proliferation signature fits the cell cycle stage requirements of infection. This preference for the differentiation signature may also be relevant to possible latency and potential reactivation in non-dividing cells (Schwartz & Stern-Ginossar, 2023). Yet, a considerable portion of the HCMV canonical ORFs (22%) have a codon usage signature that does not resemble the proliferation (r<0.6) nor the differentiation signatures (r < 0.6). They are located in the regions of the plane where very few human transcripts reside. Considering the high codon usage adaptation of human-infecting viruses to the human codon usage (Bahir et al., 2009), we expected that the fraction of lowly adapted HCMV ORFs would be smaller than we found. We further inspect the so-called “non-canonical” HCMV ORFs. Some 30% of these virus-coding genes are considered “non-canonical” is that they are only discovered through ribosome profiling (Stern-Ginossar et al., 2012). We observe here that these genes are characterized by low correlation to any of the two codon usage signatures (Figure 3B, differentiation: r < 0.6; proliferation: r < 0.6). We also found that several non-canonical genes with codon usage corresponding to the differentiation codon-usage signature are also extremely long (Figure S3B). This observation implies that long non-canonical genes are evolutionarily adapted to optimize translation, potentially indicating their role during infection.

We then continued to explore the change in the tRNA adaptation index (tAI) (Dos Reis et al., 2004) based on the tRNA pool expression of the infected cells at 24hpi. For each canonical HCMV ORF, we computed tAI relative to the tRNA pool and grouped them based on annotated functionality, as described by Ye et al., 2020. We found significant differences in mean tAI between functional gene categories. Genes involved in gene expression, DNA replication, and virion production generally have higher tAI than genes involved in host recognition and entry, viral tropism, and immunomodulation (Figure 3C, Figure S3C). We further discovered significant positive and negative correlations between the mean tAI of the functional gene groups and their similarity to the differentiation and the proliferation codon-usage signatures, respectively (Figures 3D-E). Specifically, HCMV genes with high tAI are mainly adapted to the differentiation codon usage signature and less adapted to the proliferation codon usage signature.

### tRNA - codon usage adaptation following SARS-CoV-2 infection

Different human viruses developed diverse strategies to increase the efficiency of their protein synthesis. We thus aimed to study another virus type, SARS-CoV-2, that unlike HCMV, utilizes host shut-off, which benefits the expression of the viral genes (Finkel, Gluck, et al., 2021). We examined the tRNA pool of SARS-CoV-2-infected Calu3 cells and compared them to the tRNA pool of these cells in the absence of viral infection. Since SARS-CoV-2 has a very fast replication cycle, with progeny release starting 8 hours post-infection, we only measure the tRNA pool at a single time point, 6hpi.

We found that, unlike HCMV-infected cells, the tRNA pool of SARS-CoV-2 infected cells resembled that of uninfected cells (Figure 4A, r=0.94, p<0.001). Only four tRNAs showed a significant expression dynamics. The two lowest expressing tRNA genes, Ile-AAT-3-1 and Trp-CCA-5-1, show a significant up-regulation following SARS-CoV-2 infection, while Ile-AAT-2-1 and Phe-GAA-2-1 is down-regulating following infection. We further compared the tRNA modification levels between uninfected and infected cells and found almost identical values, suggesting no change in the tRNA modification levels following SARS-CoV-2 infection (Figure 4B, r=0.99, p<0.001).

**Figure 4.**
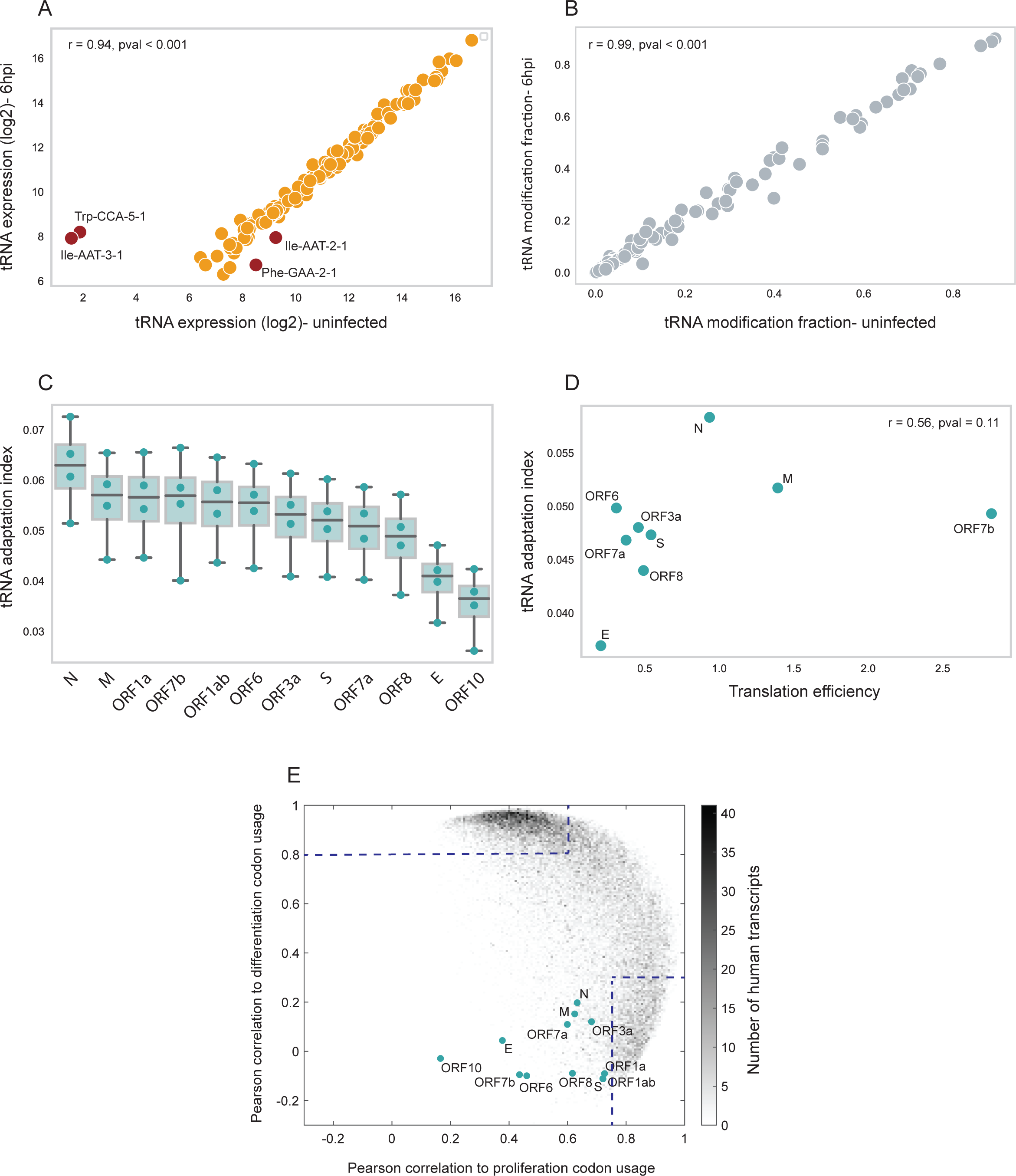
-Characterizing the tRNA pool and codon usage adaptation of SARS-CoV-2 genes following infection A| A comparison of the mean cytosolic tRNA expression level (log2) between uninfected Calu3 cells (x-axis) and 6hpi of SARS-CoV-2 infected cells (y-axis), two biological repeats. Pearson r = 0.94, p-value < 0.001. B| A comparison of the mean tRNA modification fraction between uninfected Calu3 cells (x-axis) and 6hpi of SARS-CoV-2 infected cells (y-axis), two biological repeats. Pearson r = 0.99, p-value < 0.001. C| A box plot showing the distribution of the tRNA adaptation index (tAI) values of SARS-CoV-2 genes in uninfected and infected Calu3 cells in 2 biological repeats. D| A scatter plot comparing between mean translation efficiency determined by ribosome footprint (Finkel, Mizrahi, et al., 2021) (x-axis) and mean tAI (y-axis) of SARS-CoV-2 genes. Pearson r = 0.56, p-value = 0.11. E| Density plot of the Pearson correlation measured for each SARS-Cov2 transcript to the human proliferation and differentiation codon-usage signatures. Gray dots denote human transcripts, whereas color dots denote SARS-CoV-2 genes (12 genes). The color bar (gray) represents the number of human transcripts (36762). The dashed lines represent the Pearson coefficients that determine high similarity to the proliferation or differentiation codon-usage signatures.

To discover which SARS-CoV-2 genes are adapted to the tRNA pool of their host cells, we computed the tAI for each viral gene using the tRNA expression levels in both uninfected and infected cells, as their tRNA pool is highly similar (Figure 4C). The statistical significance of differences between tAI of SARS-CoV-2 genes is shown in Figure S4A. Similar to HCMV genes, our analysis reveals that the structural genes responsible for encoding the nucleocapsid (N) and membrane (M) proteins exhibit the highest tAI values (Figure 4C), indicating a significant adaptation to the cellular tRNA pool, which likely facilitates efficient translation. Conversely, genes involved in cell entry, such as those encoding the spike (S) and envelope (E) proteins, display the lowest tAI values (Figure 4C), suggesting they are less optimized for efficient translation. Indeed, we further observed a weak positive correlation between the tAI of SARS-CoV-2 and their translation efficiency (TE) estimated by ribosome footprint (Finkel, Mizrahi, et al., 2021) (Figure 4D, r=0.56, p=0.11).

We then analyzed the codon usage adaptation of SARS-CoV-2 genes to the proliferation-differentiation codon usage signatures. Remarkably, our findings revealed a substantial alignment between the codon usage of SARS-CoV-2 genes and the proliferation rather than the differentiation signature, as depicted in Figure 4E. Notably, most SARS-CoV-2 genes diverge significantly in codon usage from human transcripts (Figure 4E). This contrasts the codon usage of HCMV ORFs, where most resemble human genes. This observation suggests that SARS-CoV-2, being a relatively new pathogen in human hosts, has not undergone extensive codon usage adaptation yet, unlike HCMV, which has been coevolving with humans since the inception of the species (Charles et al., 2023).

Similarly to HCMV, SARS-CoV-2 genes with higher similarity to the human codon usage signatures are more adapted to the tRNA pool by showing higher tAI than genes less adapted to the codon usage of human genes. This observation is mainly seen upon comparison to the proliferation codon usage signature (Figure 4E and S4B, r=0.71, p=0.009) and less so to the differentiation codon usage signature (Figure 4E and S4C, r=0.18, p=0.58).

Overall, we revealed similarities and differences in translation regulation programs during HCMV and SARS-CoV-2 infections. Both viruses show codon usage adaptation to the human codon usage signatures that manifested mainly in structural and gene expression-promoting genes. However, the tRNA pool was dynamically changing in expression and modifications only upon HCMV infection but much less so upon infection with SARS-CoV-2.

### A screen for tRNA essentiality upon cellular growth reveals essential tRNAs and a general lack of essentiality of most tRNA modification enzymes

Recently, we have manipulated the human tRNA pool with a CRISPR-based tRNA knock-out library of several tRNA families to follow their essentiality to cell proliferation and cell-cycle arrest (Aharon-Hefetz et al., 2020). We now aim to expand the scope and generate a new tRNA-CRISPR library to cover the entire repertoire of human tRNA genes, with multiple sgRNAs per tRNA gene. Here, we applied the new CRISPR library to two cellular conditions-normal cellular growth and HCMV infections.

Our newly designed CRISPR library encompasses sub-libraries targeting all types of tRNA genes, including functional cytosolic, pseudo, and mitochondrial tRNAs, along with genes encoding tRNA modification enzymes (Figure 5A). To assess the efficacy of this library under varied cellular conditions, we integrated five control sub-libraries. Among these, two sub-libraries feature dependency factors crucial for cellular growth or HCMV infection, wherein gene knock-outs result in reduced growth rates or infection efficiency, respectively. Two other sub-libraries comprise restriction factors, whereby genomic targeting enhances growth rates or infection efficiency. These factors were selected based on a previous whole-genome CRISPR library analysis of HCMV infection (Hein & Weissman, 2022). Additionally, a control group consisting of non-targeting sgRNAs was included as negative controls (Figure 5A). For a detailed description of the sgRNA library design and quality control, see supplementary file 1.

**Figure 5.**
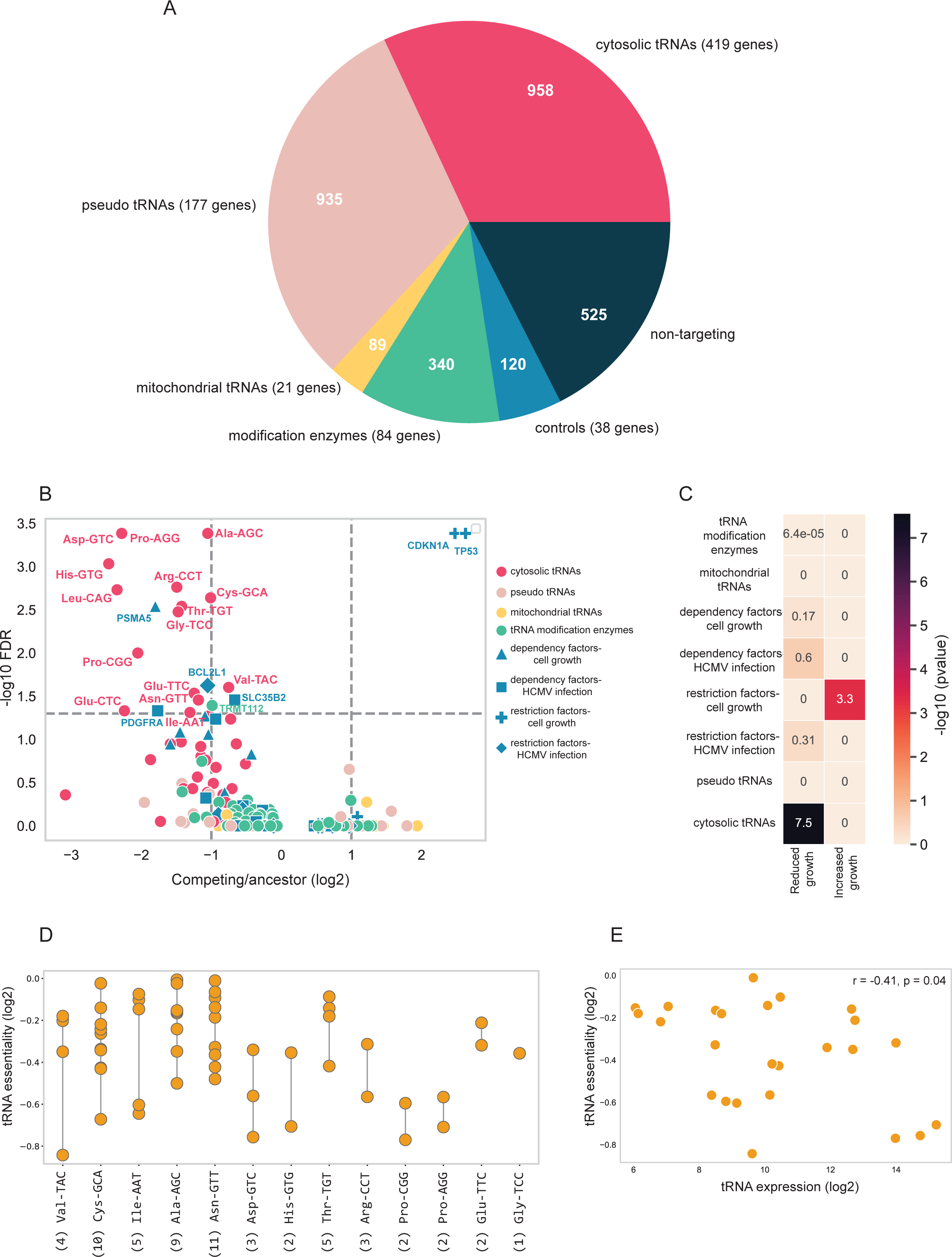
-competition experiment using tRNA-CRISPR screen revealed tRNAs that are essential for HFFs cell growth A| A pie chart describing the different sub-libraries. For each sub-library, the number of targeted genes and sgRNAs are mentioned (number of genes in brackets, number of sgRNAs within the pie chart). B| A volcano plot showing targeted gene hits from tRNA-CRISPR screen for HFFs cell growth. The X-axis shows the Z-score of log2 fold change (FC) between competing cells (3 days of competition) and ancestor samples (median of log2 FC for all high-ranked sgRNAs per gene). The y-axis shows the –log10 FDR as calculated from MAGeCK. The genes are marked according to the sub-libraries. Significance is determined by FDR < 0.05 and marked with dashed gray lines. All values are calculated for three biological repeats. C| A heat map showing the (-log10) p-value of the hypergeometric test, which tests the enrichment of each sub-library in one of the following groups: significantly reduced growth (log2FC<0, FDR <0.05) or significantly increased growth (log2FC>0, FDR <0.05). D| Differences in the essentiality of tRNA isodecoders that are the sole targets of their corresponding sgRNA. tRNA essentiality is determined by the median log2 FC of its corresponding sgRNA between competing and ancestor cells. tRNA isoaceptors are denoted on the x-axis. Each dote corresponds to a different isodecoder type within the isoaceptor family. Number of isodecoders per tRNA family is mentioned in parenthesis. E| Comparison between the (log2) expression of the tRNA isodecoder in HFFs (x-axis) and their essentiality, as shown in Figure 5D (y-axis). Pearson correlation r = -0.41, p-value = 0.04.

We initially utilize the CRISPR library’s availability to assess the essentiality of each human tRNA and modification enzyme for the cellular growth of uninfected HFFs, employing a cell competition assay. Briefly, we transduced HFFs cells with lentiviruses containing the CRISPR library in low MOI (MOI=0.3) to minimize the probability of cells being transfected with multiple sgRNAs. We thus expect that each cell will typically be targeted in one tRNA family. We note that the targeting level of the tRNA families is predicted to differ in each family due to variability in sgRNA targeting, specifically in the number of ON-targets among the tRNA isodecoders.

After two days of antibiotic selection and two days of recovery, we sampled the ancestor sample. Then, we let the cells grow for three more days and sampled the competing cell population. From sgRNA sequencing of the CRISPR-targeted cells, we determined the growth rate of each targeted cell population by comparing the fraction of each sgRNA in the cell population. To determine sgRNA and gene knock-out enrichment in competing and ancestor cells, we used the MAGeCK tool. This widely used algorithm identifies significant hits in CRISPR/Cas9 knock-out screens (Li et al., 2014). Using the MAGeCK tool, we also estimated the extent and direction of the effect of each gene hit on HFFs cell growth, i.e., increasing or reducing cell growth (Figure 5B). We further carried out a hypergeometric test to compute the significance of the overlap between the sub-libraries and their effect on growth (Figure 5C).

To validate the reliability of the CRISPR screen, we first focused on the control gene sub-groups. We observed that competing cells were significantly depleted of sgRNAs targeting several genes essential for cell growth and viability (considered as dependency factors in this screen), such as PSMA5, which controls proteolytic toxicity, and BCL2L1, which inhibits programmed cell death (Figure 5B). However, as a group, dependency factors did not significantly affect cell growth (Figure 5C). We further found that sgRNAs targeting two cell growth-restricting factors, TP53 and CDKN1A, were significantly enriched among the competing cells and also, as a group, showed a significant increase in cell growth (Figure 5B-C, p-value < 10^-3, hypergeometric test).

Continuing to explore the effect of tRNA genes knock-out on cell growth, we found that the functional cytosolic tRNA sub-library is the only tRNA gene group that significantly reduces cell growth (Figure 5C, p-value ∼=10^-7, hypergeometric test). The rest of the sub-libraries, namely pseudo-tRNAs, mitochondrial-tRNA genes, and tRNA modification enzymes, showed no effect on cell growth as entire sets (Figure 5B-C). The lack of effect on cellular growth of the pseudo-tRNAs sub-library is expected, as pseudo-tRNAs are not expressed in human cells (Figure S5A). Regarding the mitochondrial tRNA genes, as mitochondrial genes are not supposed to be targeted by the CRISPR system (Yin et al., 2022), the lack of effect of targeting these genes on cell growth (Figure 5C) is also clear. tRNA modification enzymes, as a group, do not affect growth (Figure 5B-C). Only one gene from this group, TRMT112, appears as a dependency factor for cellular growth as it was found to be depleted from the competing population upon CRISPR targeting. This gene is an activator of methyl transferases of multiple types of molecules, including tRNAs (Stelzer et al., 2016). The lack of growth effect upon targeting tRNA modification enzymes is even more surprising given that many of the tRNA modification enzymes catalyze post-transcriptional modifications on various RNA molecules besides tRNAs and are involved in multiple fundamental processes. We further compared this result by examining the phenotypic effect of the knock-outs of these enzymes in HFFs cells from a previous CRISPR screen (Hein & Weissman, 2022). In this study, tRNA modification enzymes were typically not essential (Figure S5B). One exciting possibility is that these genes fulfill crucial functions, yet other genes, e.g., paralogs, are backing them up (Kafri et al., 2006, 2008).

We further observed that for each tRNA family, different tRNA isodecoders have a different level of essentiality for cell growth (Figure 5D). We validated that the sgRNA efficiency score (as estimated from the sgRNA design tool) is not correlated with the sgRNA targeting effect on growth (data not shown). This observation resonates with previous observations that otherwise identical tRNAs might be expressed differently (Aharon-Hefetz et al., 2020; Gao et al., 2024; Sagi et al., 2016) and has a differential effect on phenotype upon knock-out (Bloom-Ackermann et al., 2014). We found a moderate negative correlation between the expression level of the targeted tRNAs in HFFs and their revealed essentiality (Figure 5E, r=-0.41, p-value=0.04). This observation suggests that highly expressed tRNA isodecoders are typically more essential for cell growth of HFFs than lowly expressed isodecoders.

### A CRISPR screen reveals tRNA genes and tRNA modification enzymes that disrupt or enhance HCMV infection upon knock-out

After screening the effect of our newly designed sgRNA library on cell growth, we set up a CRISPR screen experiment to explore the essentiality of tRNA genes and tRNA modification enzymes for HCMV infection. We applied a reporter-based CRISPR screen using an HCMV strain tagged with GFP on the UL122 (IE2) gene (Stanton et al., 2010). We transduced HFFs with the CRISPR library similarly to the previous cell transduction described above. Following antibiotic selection and two days of recovery, transduced cells were infected with IE2-GFP labeled HCMV virus in high MOI (MOI=5) to ensure a complete infection of all cells. After 72 hours, infected cells were FACS sorted based on GFP levels (Figure 6A). The sorted populations are annotated GFP1, GFP2, GFP3, and GFP4, with GFP1 corresponding to the lowly infected cells and GFP4 to the highly infected cells (Figure 6A, middle and lower panels). The GFP levels approximate the viral load in the host cell, representing the infection’s efficiency (Figure S6A). Figure 6A shows the GFP levels of non-transduced, uninfected cells (upper panel) and of transduced cells with either a single non-targeting sgRNA (as control) or the entire CRISPR library (middle and lower panels, respectively). IE2-GFP HCMV-infected cells from both the non-targeting and the library transduction exhibited strong infection, with most cells in the highest GFP bin, which specifies the strongest HCMV infection. None the less, comparing the results between these two populations revealed that in the population transduced with the library, fewer cells were found in the highest GFP bin (70% compared to 76.7%), and the three lower GFP bins (GFP1-3) were more populated (total of 28% compared to 22% in the non-targeting population). This difference indicates that some genes targeted in the library are essential for infection.

**Figure 6.**
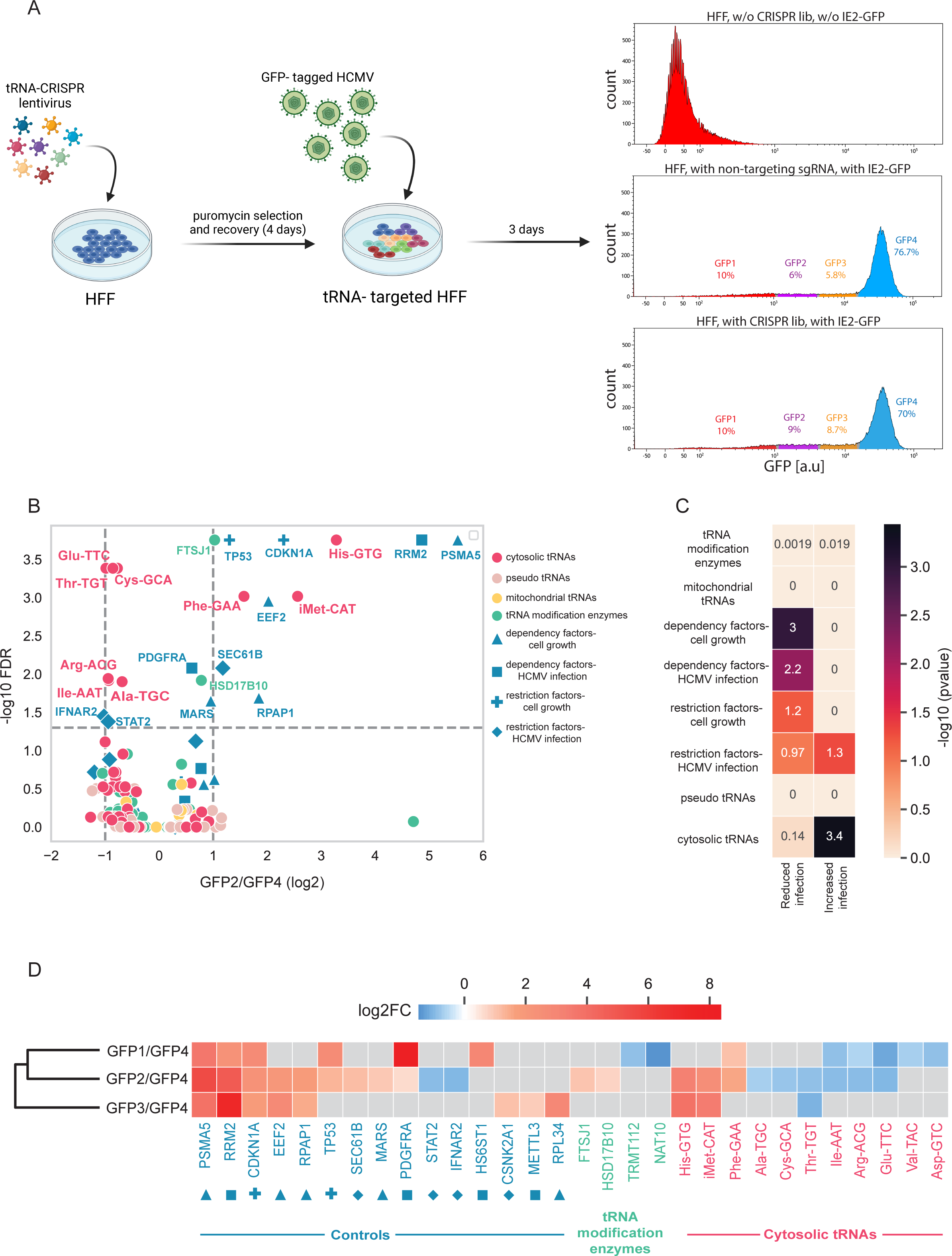
-tRNA CRISPR screen in HCMV infected-HFFs cells identified tRNA genes and modification enzymes that disrupt or improve HCMV infection upon CRISPR targeting A| A schematic representation of the experimental setup and distributions of the measured GFP levels of uninfected HFFs cells (upper panel), HCMV-infected HFFs cells with non-targeting sgRNA (middle panel), and CRISPR-targeted and HCMV-infected HFFs cells (lower panel). The GFP levels of sorted populations, GFP1-4, are marked in the middle and lower panels. B| A volcano plot for targeted gene hits from tRNA-CRISPR screen in HCMV infection. The x-axis shows the Z-score of log2 FC between lowly-infected cells (GFP2) and highly-infected cells (GFP4). The y-axis shows the –log10 FDR as calculated from MAGeCK. The genes are marked according to the sub-libraries. Significance is determined by FDR < 0.05. All values are calculated for three biological repeats. C| A heat map showing the (-log10) p-value of the hypergeometric test, which tests the enrichment of each sub-library in one of the following groups: significantly reduce infection (log2FC>0, FDR <0.05) or significantly increase infection (log2FC<0, FDR <0.05). D| A heat map describes the z-score log2FC between the different lowly-infected cell populations (GFP1, GFP2, GFP3) relative to the highly infected cells (GFP4) for the significant gene hits. Genes found as significant hits in at least one of the comparisons are presented here. Non-significant hits are marked in gray squares. The genes are colored and marked according to their sub-library, that correspond to the sub-library color and marker shape described in Figure 6B legend. The dendrogram depicts the similarity between the comparisons based on the enrichment gene hits pattern.

To estimate the effect of the variants of the sgRNA library on HCMV infection, we extracted genomic DNA from each GFP-sorted cell population and measured sgRNA abundance in each fraction by deep sequencing of the sgRNA region. We then determined the relative sgRNA enrichment between the low-GFP cell populations (i.e., GFP1, GFP2, and GFP3) and the high-GFP cell population (i.e., GFP4) using the MAGeCK tool.

The gene hits of one of the comparisons between lowly and highly infected cells, specifically GFP2 vs GFP4, are presented in Figure 6B. Reassuringly, we found that sgRNAs targeting dependency genes of HCMV infection are enriched in lowly infected cells (i.e., GFP2) (Figure 6B-C, p-value < 0.01, hypergeometric test). In contrast, sgRNAs targeting genes restricting HCMV infection are over-represented in highly infected cells (i.e., GFP4) (Figure 6B-C, p-value = 0.05, hypergeometric test). Cell growth dependency factors are also considerably enriched in lowly infected cells (i.e., GFP2) and thus reduce infection efficiency (Figure 6C, p-value = 0.001, hypergeometric test), suggesting sensitivity to the host cells’ viability and proliferation state during HCMV infection (Bogdanow et al., 2021; Fortunato et al., 2002).

After confirming the CRISPR library’s efficacy in HCMV infection by assessing dependency and restriction factors, we delved into the essentiality of tRNA families and modification enzymes. Certain cytosolic tRNAs, like His-GTG tRNA, exhibited significant enrichment in lowly infected cells (GFP2) compared to highly infected cells (GFP4), indicating their crucial role in infection by reducing infectivity post-CRISPR targeting (Figure 6B). However, collectively, cytosolic tRNA knock-outs didn’t substantially diminish infection (Figure 6C, p-value = 0.73, hypergeometric test). Conversely, certain cytosolic tRNAs acted as restriction factors, enhancing infectivity upon CRISPR targeting, exemplified by Arg-ACG tRNA (Figure 6B), with a similar collective effect (Figure 6C, p-value < 0.001, hypergeometric test). Remarkably, tRNA modification enzymes did not influence HCMV infection (Figure 6C).

Similar to previous results (Figure 5E), highly expressed tRNA isodecoders in infected cells are moderately more essential for infection, as their sgRNAs are enriched in lowly infected cells (Figure S6B). Yet, this correlation is not statistically significant (Figure S6B).

Using GFP as a reporter for the progress of HCMV infection allowed us to detect genes whose sgRNAs consistently show enhanced or reduced infection. Moreover, we detected a gradual increase in HCMV genomic DNA as GFP intensified (Figure S6A), which enabled the elucidation of delicate dynamics in gene essentiality that act at various time points along HCMV infection. We thus compared the sgRNA enrichment between all three populations of lowly infected cells, i.e., GFP1, GFP2, and GFP3, relative to the highly infected cell population, i.e., GFP4 (Figure 6D). Here, we show gene hits that are significantly enriched or depleted in at least one of the enrichment analyses. Hierarchical clustering of the comparisons shows higher similarity between GFP1 and GFP2 than GFP3 (all relative to GFP4), emphasizing that similar genes are essential in the earlier phases of infection.

To explore the temporal and directionality in the essentiality of tRNAs and tRNA modification enzymes, we compared the enrichment of their sgRNAs in each GFP-sorted population to the sgRNAs of the known dependency and restriction factors in the library. For example, sgRNA targeting Phe-GAA are highly enriched in GFP1 and GFP2 relative to GFP4, similar to PDGFRA, which is the cell receptor that enables viral entry into fibroblasts (Soroceanu et al., 2008; Wu et al., 2018) (Figure 6D). This result suggests that Phe-GAA is needed in earlier phases of infection. In contrast, sgRNAs targeting His-GTG and iMet-CAT are primarily enriched in GFP2 and GFP3, similar to the sgRNAs targeting RRM2 and METTL3 (Figure 6D). RRM2 is an essential gene for viral DNA replication (Bagga & Bouchard, 2014), and METTL3 was shown to affect viral replication as its knock-out increases IFN response, resulting in inefficient HCMV infection (Winkler et al., 2019). Thus, His-GTG and iMet-CAT tRNAs are considered dependency factors essential in later infection stages. We also observed multiple tRNAs that their sgRNAs were enriched in the highly infected cells relative to the lowly infected cell populations, such as Glu-TTC, Arg-ACG, Thr-TGT, Ile-AAT, etc. The enrichment pattern of these tRNAs hits resembles the enrichment of two known restriction factors important for IFN response, STAT2, and IFNAR2, in highly infected cells (Figure 6D), suggesting that these tRNAs are restriction factors of HCMV infection. Similar to cell growth experiments, sgRNAs targeting pseudo and mitochondrial tRNAs were not statistically enriched or depleted from any lowly infected cell populations.

sgRNAs targeting some tRNA modification enzymes showed effects on infection. sgRNAs of FTSJ1 and HSD17B10 reduced the infectivity of HCMV, as these gene hits are enriched in GFP2 cells. However, sgRNAs of TRMT112 and NAT10 were significantly depleted from GFP1, suggesting that these modification enzymes increase infectivity upon CRISPR targeting (Figure 6D). It is important to note that many of the tRNA modification enzymes, like HSD17B10, TRMT112, and NAT10, are involved in various processes, and they modify other RNA molecules except tRNAs. Therefore, it is hard to determine from the CRISPR screen which biological activity of the modification enzyme is vital for HCMV infection. FTSJ1, on the other hand, is a methyltransferase that methylates several tRNAs in the anticodon loop at positions 32 and 34 (Safran M et al., 2021) and has no other known functions. Modifications in the anticodon loop are essential for the base-pairing of the anticodon to the codon on the mRNA. Manipulation of the modification level in the anticodon loop might affect the translation accuracy of the ribosome and cause ribosomal frameshifts (Bartok et al., 2021; Pan, 2018). Enrichment of sgRNA targeting FTSJ1 in lowly-infected cells suggests that knock-out of this tRNA modification enzyme decreases the efficiency of HCMV infection, potentially through reduction of modification level on tRNAs that are involved in the translation of HCMV genes or host genes that are needed for infection.

## Discussion

In this study, we investigated the impact of tRNAs and codon usage on viral infection, focusing on HCMV and SARS-CoV-2. tRNA sequencing revealed subtle changes in tRNA expression and modifications during HCMV infection, contrasting with stable tRNA pools in SARS-CoV-2-infected cells. The codon usage of HCMV genes correlated with differentiation codon usage, while the codon usage of SARS-CoV-2 genes resembled proliferation codon usage. Structural and gene expression-related genes exhibited high adaptation to the tRNA pool in both viruses, while genes related to viral entry to cells showed lower adaptation. Applying a comprehensive CRISPR screen that targets human tRNA genes and tRNA modification enzymes, we were able to pinpoint potential tRNAs that restrict or promote HCMV infection.

Characterizing the tRNA pool of cells infected with HCMV and SARS-CoV-2 revealed the varying impact on tRNA pools in infected cells. HCMV infection leads to dynamic changes in tRNA levels in HFFs cells, while Calu3 cells infected by SARS-CoV-2 maintain a stable tRNA pool. This difference may stem from how SARS-CoV-2 induces host shut-off (Finkel, Gluck, et al., 2021), reducing host mRNA levels and potentially minimizing the need for tRNA adjustments. In contrast, HCMV employs strategies that rely on continued host translation machinery (McKinney et al., 2014; Tirosh et al., 2015), prompting manipulation of the tRNA pool to potentially benefit viral infection. Previous studies have identified various tRNA isodecoders regulated in different organisms and tissues (Barski et al., 2010; Dittmar et al., 2006; Rak et al., 2021; Sagi et al., 2016; Torrent et al., 2018; Z. Zhang et al., 2018). Here, we also found a wide range of expression changes in tRNA isodecoders following HCMV infection, with some genes positively or negatively regulated. CRISPR screening supported this observation, indicating that highly expressed tRNA isodecoders substantially impact cell growth and HCMV infection more than lowly expressed ones. Reduction in dihydrouridine modification was observed in three tRNAs post-HCMV infection, potentially contributing to their downregulation (Faivre et al., 2021). Further validation is required to confirm the causal effect of this modification on tRNA stability in HCMV infection.

Sequencing tRNA pools of IFN-treated and HCMV-infected cells showed a minor effect of the IFN response, suggesting that the virus, as opposed to the cell, is the agent that primarily drives changes in the tRNA pool. However, instances where the host response opposed the virus’s impact on tRNA expression and modification levels were observed, highlighting the complexity of the interplay. For example, Ser-AGA tRNA was downregulated in IFN-treated cells but up-regulated in late HCMV infection. The modification level of ms2t6A on Lys-TTT tRNAs increased in IFN-treated cells but gradually decreased during HCMV infection. However, the potential regulators responsible for these changes remain unknown.

We found certain changes in the level of modifications in the anticodon loop following infection, specifically ms2t6A on Lys-TTT and m3C on Ser-TGA. Several publications recently identified changes in the modification level of wybutosine on Phe-GAA and ms2t6A on Lys-TTT in proliferating cancerous cells (Rosselló-Tortella et al., 2020; Smith et al., 1985) and in T-cells that proliferate upon antigen-induced activation (Rak et al., 2021). It was also shown that m3C modification on Ser-TGA is associated with the pluripotency of stem cells and tumor cell growth (Ignatova et al., 2020). Overall, these two types of modifications correlate with cell proliferation. HCMV infection is known to be most efficient in infecting quiescent cells, which restart their cell cycle upon infection and are then arrested before entering into the S-phase (Bogdanow et al., 2021; Fortunato et al., 2002). In this phase, the virus exploits the massive production of essential raw materials for viral DNA replication, like dNTPs, made by the recycling cell (Lembo et al., 1999; Gupta & Mlcochova, 2022). The change in these modifications suggests that they are related to the cycling phase of the cells, a crucial step for a proper HCMV infection.

To explore if changes in tRNA levels post-viral infection enhance viral gene translation based on their codon usage, we compared viral and human genes’ codon usage. While focusing on proliferation and differentiation states, this comparison revealed HCMV genes correlating with differentiation codon usage and SARS-CoV-2 genes with proliferation codon usage. This result aligns with the vast diversity of host cells they infect, highlighting tissue tropism as a critical determinant of viral codon usage (Hernandez-Alias et al., 2021). A comprehensive comparison between the codon usage of viral proteins and the dichotomous codon usage signatures of human tissues can explain codon usage adaptation of human-infecting viruses to the cell type they infect, constituting a significant factor in shaping the genomes of the evolved human-infecting viruses. Previous studies comparing codon usage adaptation among human infecting viruses and other viruses have shown a significant correlation between the codon usage of genes of human viruses and their hosts. However, our research reveals a noteworthy proportion of viral genes exhibiting a low correlation with proliferation or differentiation codon-usage patterns, which holds even for HCMV genes adapted to infect human cells (Charles et al., 2023). However, HCMV genes may evolve to other codon usage signatures of human genes, like stress-related genes, that we did not explore here. Further characterization of low-compatibility genes to the proliferation-differentiation codon usage signatures is necessary to comprehensively understand viral codon usage adaptations.

To assess tRNA essentiality in HCMV infection, we designed the first comprehensive CRISPR library targeting all tRNA modification enzymes and human tRNA genes, including functional cytosolic tRNAs and pseudogenes. The library consists of multiple sgRNAs per tRNA gene, and each sgRNA potentially targets multiple tRNA genes belonging to the same tRNA family in high precision. Targeting the same tRNA gene with various sgRNAs, each having a different targeting site, enabled robust and reproducible results. This systematic approach allowed for the first time to knock-out each tRNA family and modification enzymes, ensuring maximal impact on the translational machinery. Here, we examine the newly designed CRISPR library in the context of cellular growth and viral infection. However, this library can be applied to various cellular conditions and contexts. We found that 15 tRNA families, out of 49 tRNA families, reduced cell fitness during the proliferation of HFFs, highlighting their essential role. While our previous study (Aharon-Hefetz et al., 2020) showed higher essentiality of proliferation-related tRNAs in HeLa cells, this pattern did not hold in slowly dividing WI38 cells. In agreement, in the current work, we found that HFFs growth depends on both proliferating and differentiation-related tRNAs. This supports the conclusion that slowly dividing cells rely on both types of tRNAs for cellular processes, such as cell division and multicellularity.

We used a GFP reporter in the HCMV infection CRISPR screen to monitor the dependency of infectivity stages on tRNA families and tRNA modification enzymes. sgRNAs enriched or depleted in lowly infected cells indicate genes essential or restrictive to HCMV infection, respectively. One of the gene hits that its corresponding sgRNAs were enriched in lowly infected cells was tRNA His-GTG. Intriguingly, this tRNA is selectively encapsulated in HCMV virions and was suggested to contribute to stabilizing the nucleocapsid to support the large dsDNA genome of HCMV (Liu et al. 2021). Our results indicate that His-GTG might have an additional role during HCMV infection, probably at the late stages of infection. We further found that cells with deletion of tRNA that encodes the initiator methionine (iMet-CAT) were enriched in lowly infected cells. Interestingly, it was previously shown that up-regulation of iMet-CAT modulates the tRNA pool, increases cell proliferation, and promotes tumorigenesis (Clarke et al., 2016; Pavon-Eternod, Gomes et al., 2013). iMet-CAT may be essential for HCMV infection by affecting viral-mRNA translation or via affecting the modulation of the cell cycle by the virus (Tucker et al., 2020).

Intriguingly, we found several tRNAs depleted from lowly infected cells, implying that their regular presence in cells restricts HCMV infection. These tRNAs might limit infection by interaction with the anti-viral stress response. For example, SLFN proteins, such as SFLN13 and SFLN8, cleave tRNAs in their 3’ end, an activity that reduces the ready-to-translate tRNAs and the production of viral particles (Yang et al., 2018). Previous studies showed that lowering tRNA expression improves protein folding by decreasing translation elongation speed (Gorochowski et al., 2015). Additional studies are needed to understand better how these tRNAs restrict or are required for HCMV infection.

tRNA modification enzymes are vital for tRNAs and sometimes other RNAs maturation, and they participate in additional cellular processes. Thus, we anticipated that they would constitute significant hits in our CRISPR screens as a group. Surprisingly, these enzymes were mainly found to have little effect on cell proliferation or HCMV infection. Two hypotheses could rationalize this observation. One is inefficient targeting by the CRISPR system due to inefficient sgRNAs, limited accessibility of the gene’s loci to the Cas9, etc. (Horlbeck et al., 2016; Yuen et al., 2017). Another option is gene redundancy, as previous studies showed for genes with crucial functions (Kafri et al., 2006, 2008), which renders them non-essential despite their essential function. Further studies on the potential backup mechanism between tRNA modification enzymes are necessary to understand their role in viral infection.

In conclusion, our investigation sheds light on the intricate interplay between tRNAs, codon usage, and viral infection, particularly focusing on HCMV and SARS-CoV-2. These findings contribute to a deeper understanding of the molecular mechanisms underlying viral-host interactions and offer new avenues for future research in antiviral strategies and therapeutic development.

## Materials and methods

### Cell culture

HEK293T cells (ATCC; CRL-3216) and Human Foreskin Fibroblasts (ATCC; CRL-1634) were grown in DMEM high glucose medium (Biological Industries; 01-052-1A) supplemented with 10% heat-inactivated FBS, 1% penicillin/ streptomycin (P/S) and 1% L-Glutamine.

Calu3 cells (ATCC HTB-55) were cultured in 6-well or 10-cm plates with RPMI supplemented with 10% fetal bovine serum (FBS), MEM non-essential amino acids, 2 mM L-glutamine, 100 units per ml penicillin and 1% Na-pyruvate.

### Mature tRNA sequencing of viral-infected cells

HFFS cells (passage 27) were grown to full confluence in 10 cm plates and then infected with the HCMV merlin UL32-GFP strain (Stanton *et al*. 2010) with an MOI of 5. Cells were incubated with the virus for 1 hour, washed, and supplemented with fresh medium. Cells were harvested at 4-time points following infection-5, 16, 24, and 72 hours post-infection. HFFs treated with IFN were incubated with 550U/ml IFNα (PBL; 11200-2) and 700U/mL IFNβ (Peprotech; 300-02BC-20) and harvested after 5 hours. Infected cells treated with ruxolitinib were supplemented with a medium containing 4uM ruxolitinib (InvivoGen; 941678-49-5) after 1 hour incubation with HCMV and harvested 24 hours post-infection.

SARS-CoV-2 infection of Calu3 cells and RNA extraction was done as described in (Finkel, Mizrahi, et al., 2021). Sequencing library preparation and data analysis were done as described in (Rak et al., 2021).

### Codon usage analysis and tAI estimation from tRNA expression

Estimating tAI from viral-infected cells was done as described in (Rak et al., 2021).

### sgRNA library design

The sequences of all human tRNA genes, including functional cytosolic, pseudo and mitochondrial tRNAs, were downloaded from the tRNA database (Chan and Lowe 2016). The CRISPR sgRNA design tool of Benchling ([Biology Software]. (2022). Retrieved from https://benchling.com), was used for sgRNA design, using the default parameters. To remove sgRNAs with tRNA off-targets (sgRNAs that have high sequence similarity to tRNAs which are not part of the targeted tRNA family), we used the Blast tool (Altschul *et al*. 1990) and compared each sgRNA sequence (with a pam sequence NGG downstream) to the entire tRNA gene sequences (both functional cytosolic and pseudo tRNAs). sgRNAs with 0-1 mismatches relative to a non-targeted tRNA were removed from the final list of sgRNAs, provided the targeted tRNAs have at least two valid sgRNAs.

### sgRNA library cloning

The sgRNA library was synthesized as a DNA oligo pool by Twist Biosciences. In addition to the sgRNA sequence, all oligos contained flanking regions containing several elements, including restriction sites for cloning and primers for sub-library amplification. Oligos were designed using the CRISPR clue tool (Becker et al., 2020). The final oligos design is shown in supplementary Figure S7H, and their sequences are shown in supplementary data 1.

The synthesized oligos were amplified by PCR, using 2.5ng of oligos, 10uM of general primers, and X2 Kapa HiFi ready mix [Roche; KK2602], with a total volume of 50ul. The Tm of the PCR was 55^0^C, and 14 cycles were done to amplify the sgRNAs. The PCR product was cleaned using a PCR clean-up kit (Promega, A9281). The PCR product was then cloned into a lentiviral vector using Golden Gate cloning (Benchling, [Biology Software]. (2022). Retrieved from https://benchling.com). Briefly, 50ng of LentiCRISPR v2 (addgen, #52961) and 5ng of PCR-amplified sgRNA pool was mixed with 1ul of BsmB1-V2 restriction enzyme (NEB; #R0580) and 1ul of T7 ligase (NEB; M0202). Digestion and ligation of the vector to the oligos were conducted by 50 cycles of changing temperature (42.5-16°C, 5 minutes each). The mixture was heated to 60°C for 5 minutes to terminate the digestion reaction. Then, the cloned vector was cleaned using standard ethanol precipitation (Sambrook and Russell 2001) and electroporated into 50ul of NEB stable *E.coli* strain (NEB; C3040H) using 1ul of cloning material (Applied volts-2200V, Resistance-200, Capacitance-25uF). Transformed cells were recovered in SOC media for an hour at 37°C, then selected by growing on LB medium supplemented with 100ug/ml ampicillin for 16 hours at 25°C. Transformation yielded X800 library coverage, as determined by cell seeding on selection plates in serial dilution. Finally, cells were harvested at OD=1 for plasmid extraction using the NucleoBond Xtra Midi kit (Macherey-Nagel; 740412.50). The composition of the final library was verified by deep sequencing.

### Cell transduction with lentiviral-tRNA-CRISPR library and HCMV infection

HEK293T cells were seeded onto 10 cm plates so that cell confluence would be approximately 70% the next day. A day after, 2.5µg of PMD2.G (Addgene; 12259) and 2.5µg psPAX2 (addgene;12260) packaging vectors were co-transfected with 5µg of the sgRNA library using 30µl of jetPEI (Polyplus; 101-10N) in DMEM high glucose medium (10ml). After 60 hours, the lentivirus-containing medium was collected and centrifuged for 15 minutes at 3200g, 4°C. The supernatant was collected in a new tube and filtered with a 0.4nm filter. The lentivirus-containing media was stored in aliquots at -80°C. The cell death curve of HFFs determined the viral titer. HFFs (passage 25) were transduced at a low MOI (0.3) to ensure that most cells receive only one viral construct with a high probability, 8ug/ml of Hexadimethrine Bromide (Sigma Aldrich; H9268). 48 hours after transduction, 1.75ug/ml puromycin (Thermo Fisher; A1113802) was added to select for infected cells. Then, the media was refreshed (without antibiotics), and cells were recovered for 48 hours. Six days after lentiviral transduction, the cells were infected with the HCMV Merlin strain with a GFP tag fused to the IE2 gene (Stanton et al. 2010), with a high MOI of 5 as follow: cells were incubated with the virus for 1 h, washed, and supplemented with fresh DMEM medium. 72 hours post-HCMV infection, cells were harvested in a medium enriched with 2% FCS, and FACS sorted into 4 cell populations based on GFP intensity (GFP1 to 4). Flow cytometry analysis was performed on a BD FACSAria Fusion instrument (BD Immunocytometry Systems) equipped with 488-, 405-, 561-and 640-nm lasers, using a 100-μm nozzle, controlled by BD FACS Diva software v8.0.1 (BD Biosciences). GFP was detected by excitation at 488 nm and collection of emission using 502 longpass (LP) and 530/30 bandpass (BP) filters. Each subpopulation was centrifuged after sorting, and cell pellets were stored at -80°C. To identify sgRNAs that target essential genes for cell viability, cells with the CRISPR library were grown similarly to the tested cells without HCMV infection. These experiments were done in three biological repeats.

### sgRNA sequencing-library preparation and data processing

Genomic DNA was extracted from each sorted population, ancestor samples, and samples without HCMV infection using the NK lysis protocol (Chen et al., 2015). Genomic DNA was used as a template for PCR to amplify the sgRNAs, as was described in (Aharon-Hefetz et al., 2020). Here, we used 1ug of genomic DNA for the first PCR reaction and ran the PCR program for 16 cycles. Shifted primers were used to increase library complexity. MAGeCK software quantifies and tests for sgRNA enrichment (Li et al., 2014). The abundance of sgRNAs was first determined using the MAGeCK “count” module for the raw fastq files. The MAGeCK “test” module was used with default parameters to test for robust sgRNA and gene-level enrichment.

### Quantitative real-time PCR analysis

Real-time PCR was performed on the extracted genomic DNA of the GFP-sorted cell populations using the SYBR Green PCR master-mix (ABI) on the QuantStudio 12K Flex (ABI) with the following primers (forward, reverse): UL55 (TGGGCGAGGACAACGAA, TGAGGCTGGGAAGCTGACAT); and B2M (CTCAACACGGGAAACCTCAC, CGCTCCACCAACTAAGAACG). To estimate the number of viral genomes in each GFP sorted sample, the following formula was calculated: 𝑛𝑢𝑚 𝑜𝑓 𝑣𝑖𝑟𝑎𝑙 𝑔𝑒𝑛𝑜𝑚𝑒𝑠 = 2^𝐶𝑡[𝑔]−𝐶𝑡[𝑛]^; where Ct[g] is the Ct value of UL55 gene and Ct[n] is the mean Ct value of all technical repeats of the normalizer gene B2M. Three technical replicates of one representative experiment are shown. Due to the low amount of genomic material of GFP1, qPCR analysis was not performed on this sorted cell population.

## Supplementary file 1

### Design of a comprehensive sgRNA library for the human tRNA pool

Recently, we have manipulated the human tRNA pool with a CRISPR-based tRNA knockout library of several tRNA families to follow the essentiality of these tRNA families for cell proliferation and cell cycle arrest (Aharon-Hefetz et al., 2020). Here, we aim to expand the scope and generate a new tRNA-CRISPR library to cover the entire repertoire of human tRNA genes, with multiple sgRNAs per tRNA gene. The new CRISPR library we designed comprises sub-libraries in which we targeted both tRNA and protein-coding genes (Figure 5A). The version of the human tRNA pool we worked with (Chan & Lowe, 2016) consists of 617 tRNA genes covering 49 families of functional cytosolic tRNAs, 54 pseudo tRNA gene families, and 21 mitochondrial tRNA genes, all of which we have aimed to target here (Figure 5A). We note that mitochondrial tRNAs are not supposed to be targeted by the current CRISPR/Cas9 editing method since the Cas9 enzyme functions in the cytosol. Thus, at the bare minimum, these tRNAs can be expected to serve as a neutral control for the CRISPR edits, as they are not likely to be changed. Due to the high similarity between the tRNA genes within isoacceptor, i.e., the same anticodon family (near 100% identity among about half of the tRNA genes belonging to the same isodecoder tRNA family), we could potentially target multiple tRNA isodecoder genes with the same sgRNA. Yet, the sequence similarity between different families may come with the challenge of undesired off-targeting of tRNAs from other tRNA families. To reduce the off-target effect between tRNA families, after we designed sgRNA candidates for each human tRNA gene, we filtered out the sgRNAs that had between 0 to 1 mismatches relative to other tRNAs that were not part of the targeted family, as long as the targeted tRNAs have at least two other potential sgRNAs. In our final set of sgRNAs, the number of sgRNAs per tRNA family varies significantly between tRNA isoacceptors of both functional and pseudo tRNAs (Figure S7A-B). On average, there are seven sgRNAs per functional cytosolic or pseudo tRNA gene, 19.5 sgRNAs per functional cytosolic tRNA family, 17.3 sgRNAs per pseudo tRNA family, and 4.2 sgRNAs per mitochondrial tRNA. In both the functional cytosolic and pseudo tRNAs sublibraries, the editing position of most of the sgRNAs is downstream of the anticodon (Figure S7C-D). The efficiency score of the sgRNAs, based on the sgRNA design algorithm, is highly variable (Figure S7E-F). We did not exclude low-efficiency scoring sgRNAs from the final library because some of them proved to reduce the targeted tRNA expression up to 2 fold in the previous library (Aharon-Hefetz et al., 2020).

Apart from sgRNAs targeting tRNAs, we included in this CRISPR library sgRNAs that target tRNA-related protein-coding genes (Figure 5A). In particular, the change in certain chemical modifications on tRNAs we observed during HCMV infection encouraged us to test the essentiality of tRNA modification enzymes during HCMV infection. Thus, a sub-library consisting of sgRNAs that target 84 out of 90 known genes that encode for tRNA modification enzymes (Ashburner et al., 2000; Carbon et al., 2009; Consortium et al., 2023) (Figure 5A) was added. Another sub-library consists of sgRNAs that target other human protein-coding genes and serve as known dependency and restriction factors for the HCMV infection model and cell proliferation, which we call the control group (Figure 5A). We chose the control genes based on the study of Hein *et al*., where they performed a whole genome CRISPR library screening in HCMV infection (Hein & Weissman, 2022). We chose between 5 and 8 genes for each sub-group: restricting and dependent factors for the HCMV infection and cell growth (Figure S7G). These sgRNAs are thus expected to enhance or reduce infection or cellular growth, respectivly. In addition, we added 525 non-targeting sgRNAs (Figure 5A) that consist of random nucleotide sequences, which will serve as neutral controls, as customary in such CRISPR-based knockout libraries (Doench et al., 2015). The sequences of sgRNAs we used for the two protein-coding sub-libraries and the non-targeting sgRNAs were taken from the Brunello library (Doench et al. 2015), while the sgRNAs for all human tRNAs were designed in-house. Our library consists of the first CRISPR sgRNA library aiming to target the entire human tRNA pool (see M&M).

### Testing the targeting quality of the sgRNA library

We tested the quality of the CRISPR screen experiment by several criteria. In each sample, pairs of sgRNAs that target the same gene are expected to show similar essentiality values compared to teams of sgRNAs that do not target the same genes. Figure S8A shows that sgRNAs for the same gene typically behave more similarly than sgRNAs for different genes. Using the same ratio test described above, we found that the targeting quality of our designed sgRNAs against tRNA genes is comparable to the well-defined sgRNAs targeting protein-coding genes (Doench et al., 2015) (Figure S8B). We further tested the representation of the non-targeting sgRNAs in the CRISPR-targeted population. Non-targeting sgRNAs serve as negative controls, and we indeed observe that they are not significantly enriched in the CRISPR-targeted cells following competition (Figure S8C). Next, we examined the quality of the highly-ranked sgRNAs, as determined by the MAGeCK tool. We found that high-ranked sgRNAs target tRNA genes with higher expression levels in HFF cells than those targeted by low-ranked sgRNAs (Figure S8D). This result suggests that the high-ranked sgRNAs, which contribute to the phenotypic effect, target highly expressed tRNAs within the cellular tRNA pool.

**Figure S1.**
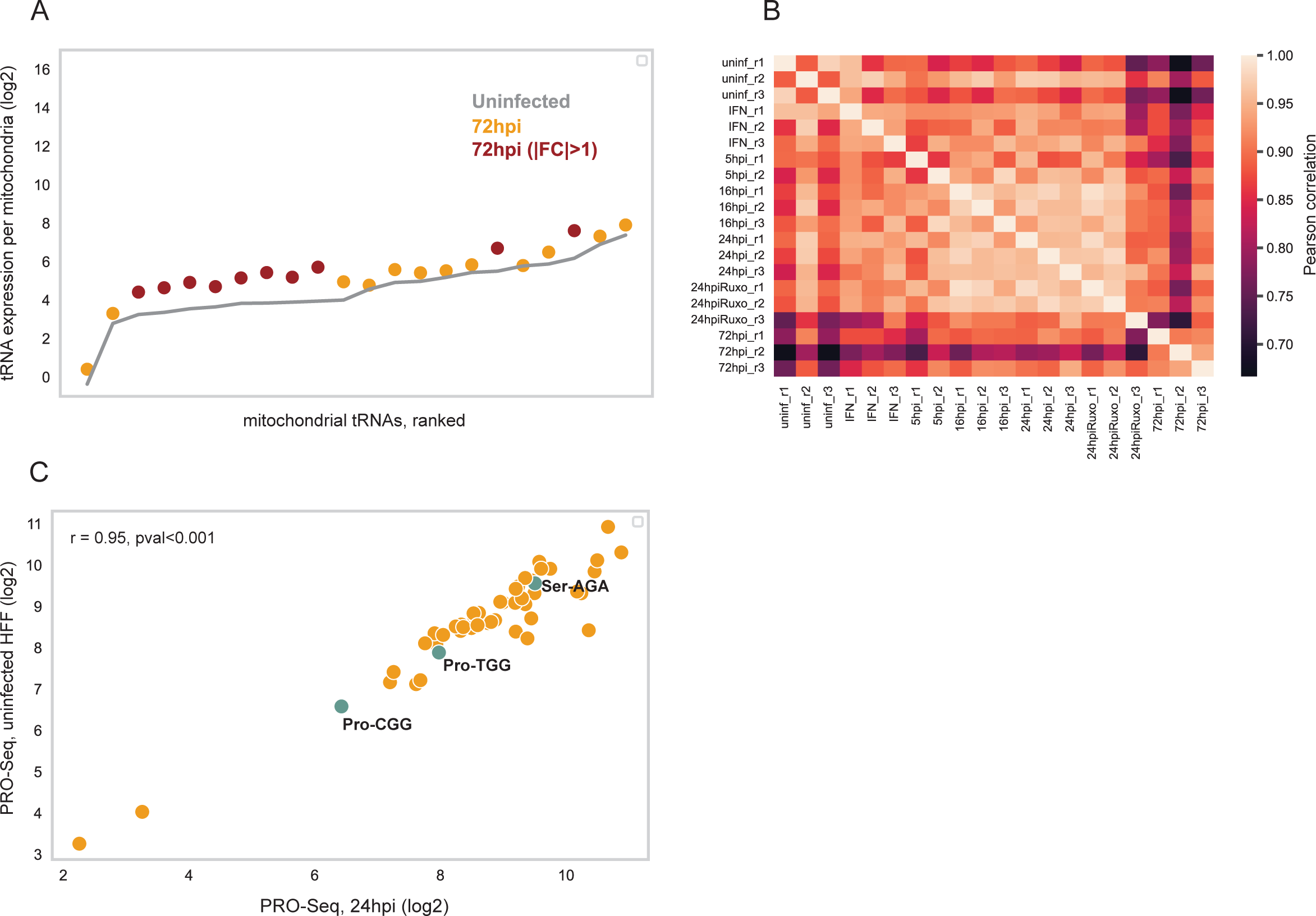
-A| Mitochondrial tRNA expression (log2) of uninfected HFFs (in gray line) and HCMV-infected cells (in yellow-red) at 72hpi normalized to the typical number of mitochondria per cell. The mitochondrial tRNAs are ordered based on their expression level in uninfected cells. tRNA genes that are differentially expressed (|FC|>1) are marked in dark red. B| Pearson correlation matrix between tRNA expression of all samples and biological repeats. C| A comparison of nascent tRNA expression levels between HCMV infected cells at 24hpi (x-axis, log2) and uninfected HFFs (y-axis, log2). The data produced by PRO-SEQ technology was taken from (Ball et al., 2022). Each dot represents the sum expression of all tRNA isodecoders belonging to the same tRNA isoacceptor. The marked tRNA isoacceptors in green refer to the differentially expressed tRNAs that show opposing dynamics between IFN-treated and HCMV-infected cells (Figure 1F). Pearson correlation r=0.95, p-value<0.001.

**Figure S2.**
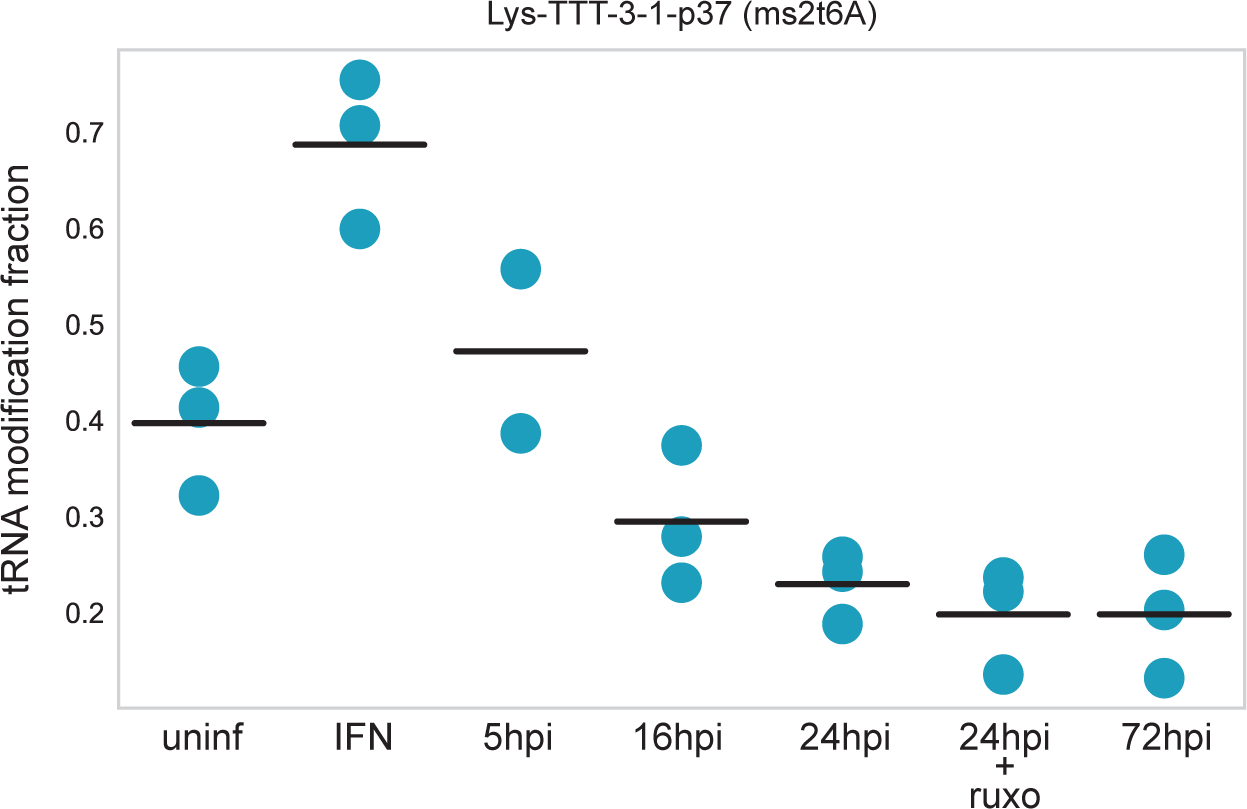
-Change in the tRNA modification level along HCMV infection on Lys-TTT-5-1 gene, position 37, modification ms2t6A. For each sample, each dot depicts a biological repeat (3 repeats in total). The line represents the average modification level in each condition or HCMV infection time point. Shown here is the same modification at the same position and same tRNA type as in Figure 2B, yet on a different tRNA isodecoder.

**Figure S3.**
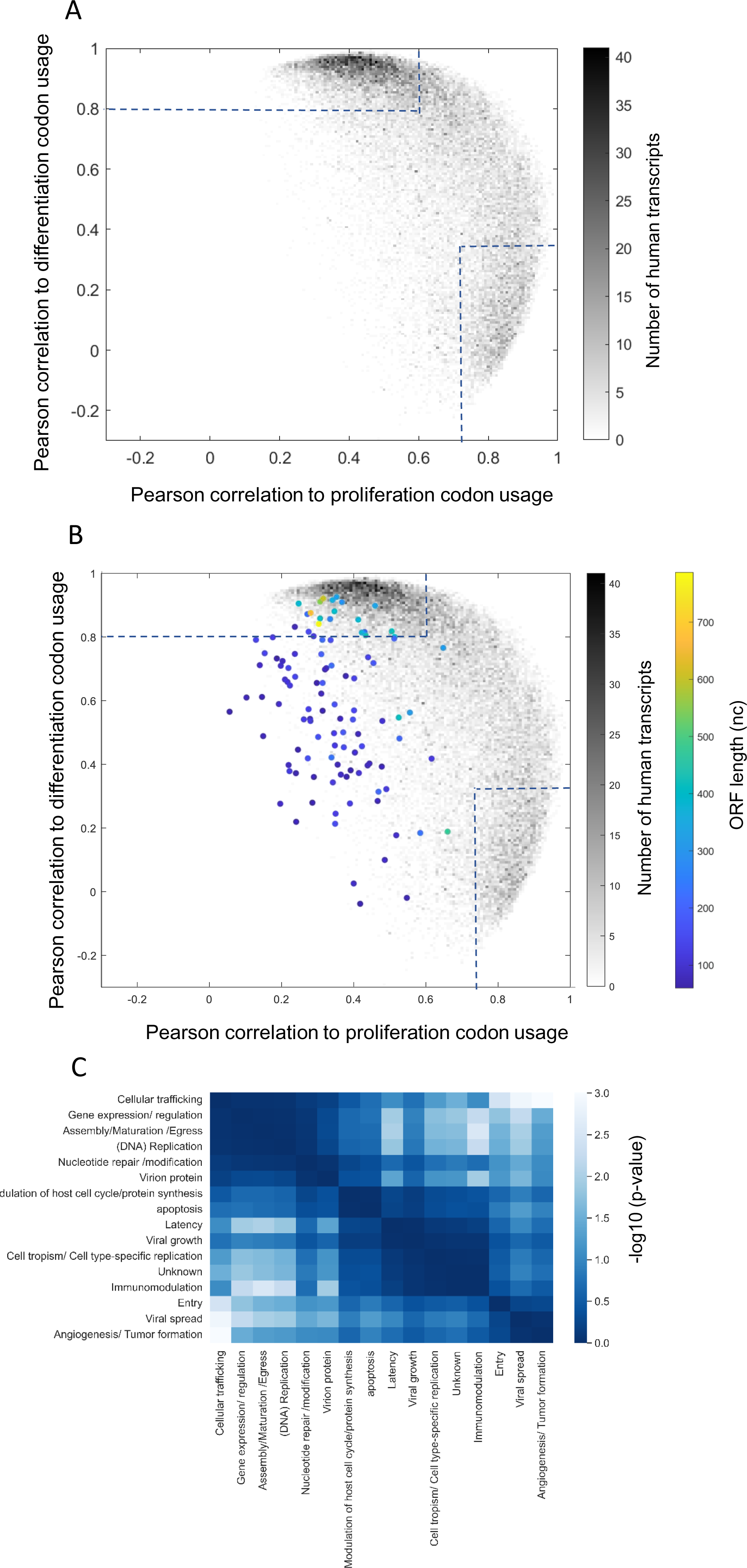
-A-B| Density plots of the Pearson correlation measured for each transcript to the human proliferation and differentiation codon-usage signatures. Gray dots denote human transcripts. The color bar (gray) represents the number of human transcripts (36762). The dashed lines represent the Pearson coefficients that determine high similarity to the proliferation or differentiation codon usage signatures. In B| color dots denote non-canonical HCMV ORFs. The color bar of the non-canonical HCMV ORFs depicts the nucleotide length of the gene. The analysis did not include short non-canonical ORFs (codon count < 58). C| A heatmap describing the (-log10) p-values of pairwise T-test that test for significant differences in the tAI levels between functional gene groups of HCMV.

**Figure S4.**
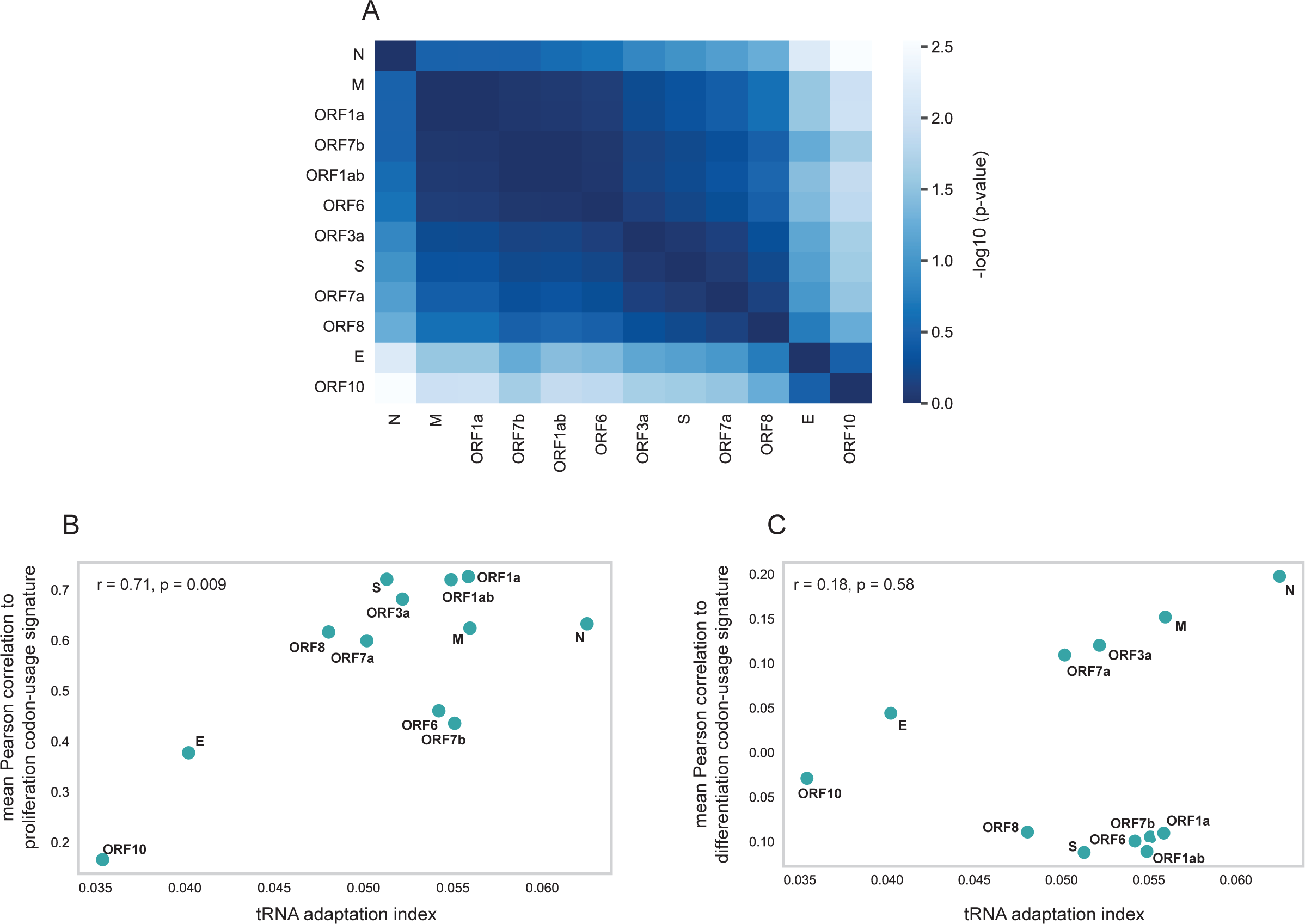
-A| A heatmap describing the (-log10) p-values of pairwise T-test that test for significant differences in the tAI levels between SARS-CoV-2 genes. B-C| Scatter plots comparing between tAI of SARS-CoV-2 genes (x-axis) and their Pearson correlation to B| proliferation, C| differentiation codon-usage (y-axis). The measured tAI is averaged tAI calculated from the tRNA pool of uninfected Calu3 cells and SARS-CoV-2 infected cells in two biological repeats.

**Figure S5.**
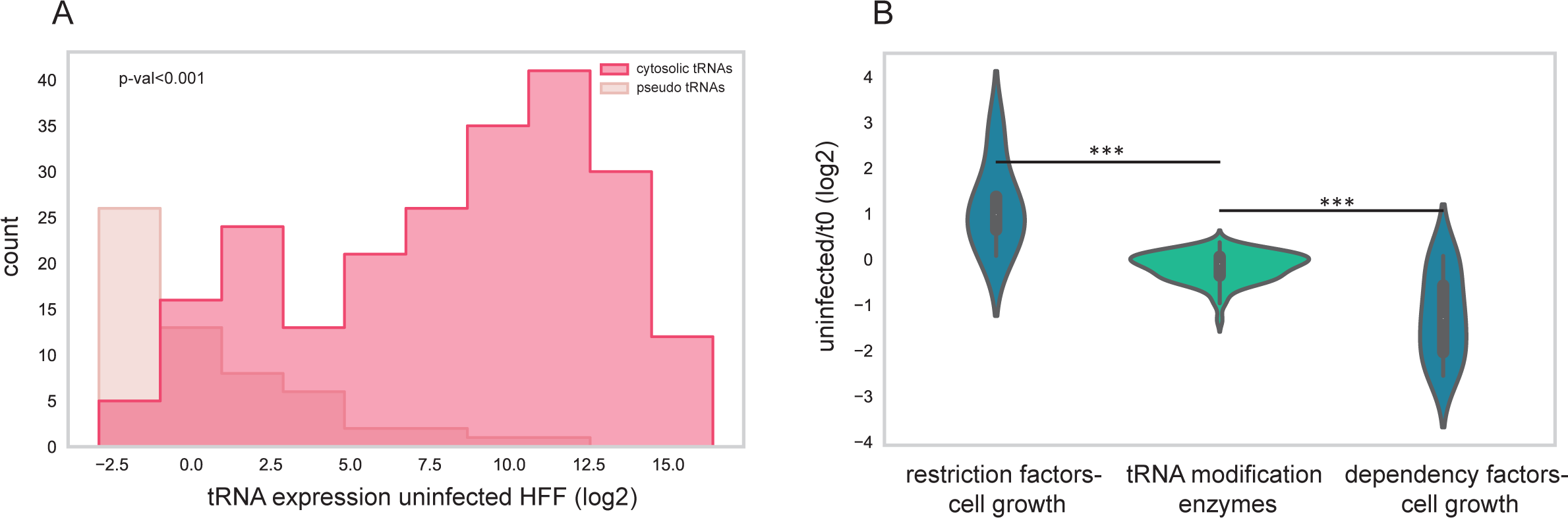
-A| Histograms describing the tRNA expression (log2) of functional cytosolic tRNAs (red) and pseudo tRNAs (beige). B| Violin plots comparing the enrichment of sgRNAs in competing cells relative to the ancestor cells taken from (Hein & Weissman, 2022) that target three sub-libraries: restricting factors for cell growth, tRNA modification enzymes, and dependency factors for cell growth. T-test resulted in a significant difference (p-value<0.001) in sgRNA enrichment between sgRNAs targeting tRNA modification enzymes and sgRNAs targeting the control sub-libraries.

**Figure S6.**
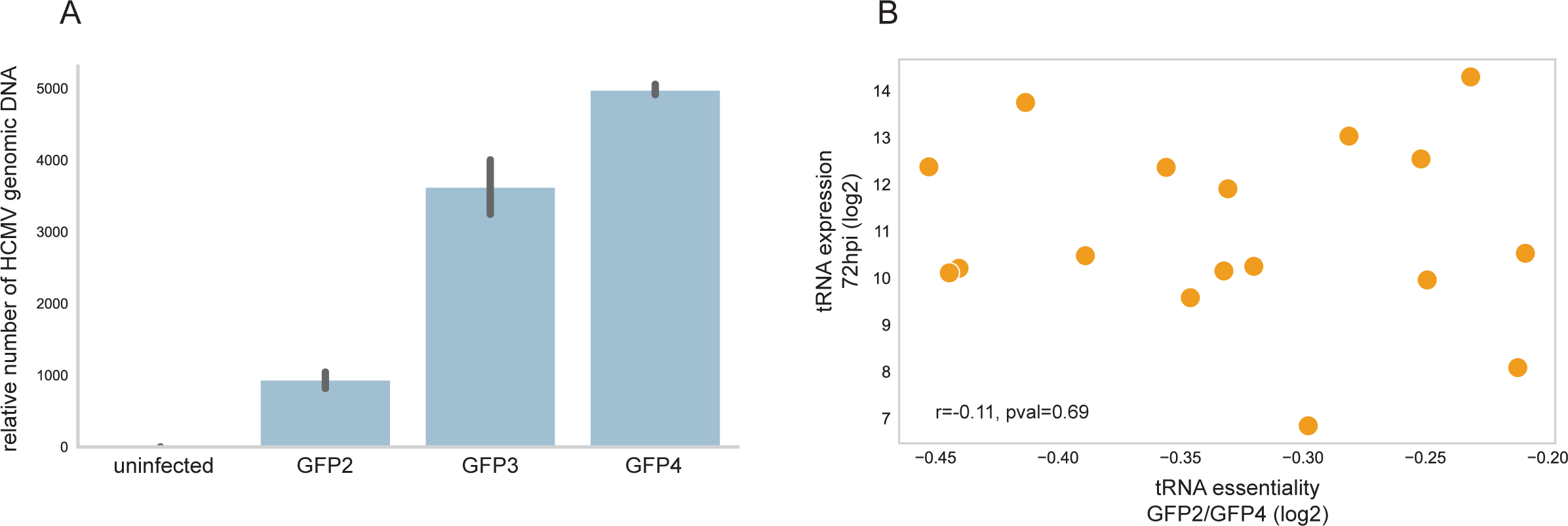
A| Number of viral genomes estimated by the relative number of UL55 normalized to B2M human gene, calculated by qPCR, in each GFP sorted cell population. The error bars depict three technical repeats. B| Comparison between the essentiality of the tRNA isodecoder to HCMV infection (x-axis) as determined by the log2 FC of its sgRNA between GFP2 and GFP4 sorted cells and the (log2) expression of the corresponding tRNAs in infected cells, 72hpi (y-axis). Pearson correlation r = - 0.11, p-value = 0.69.

**Figure S7.**
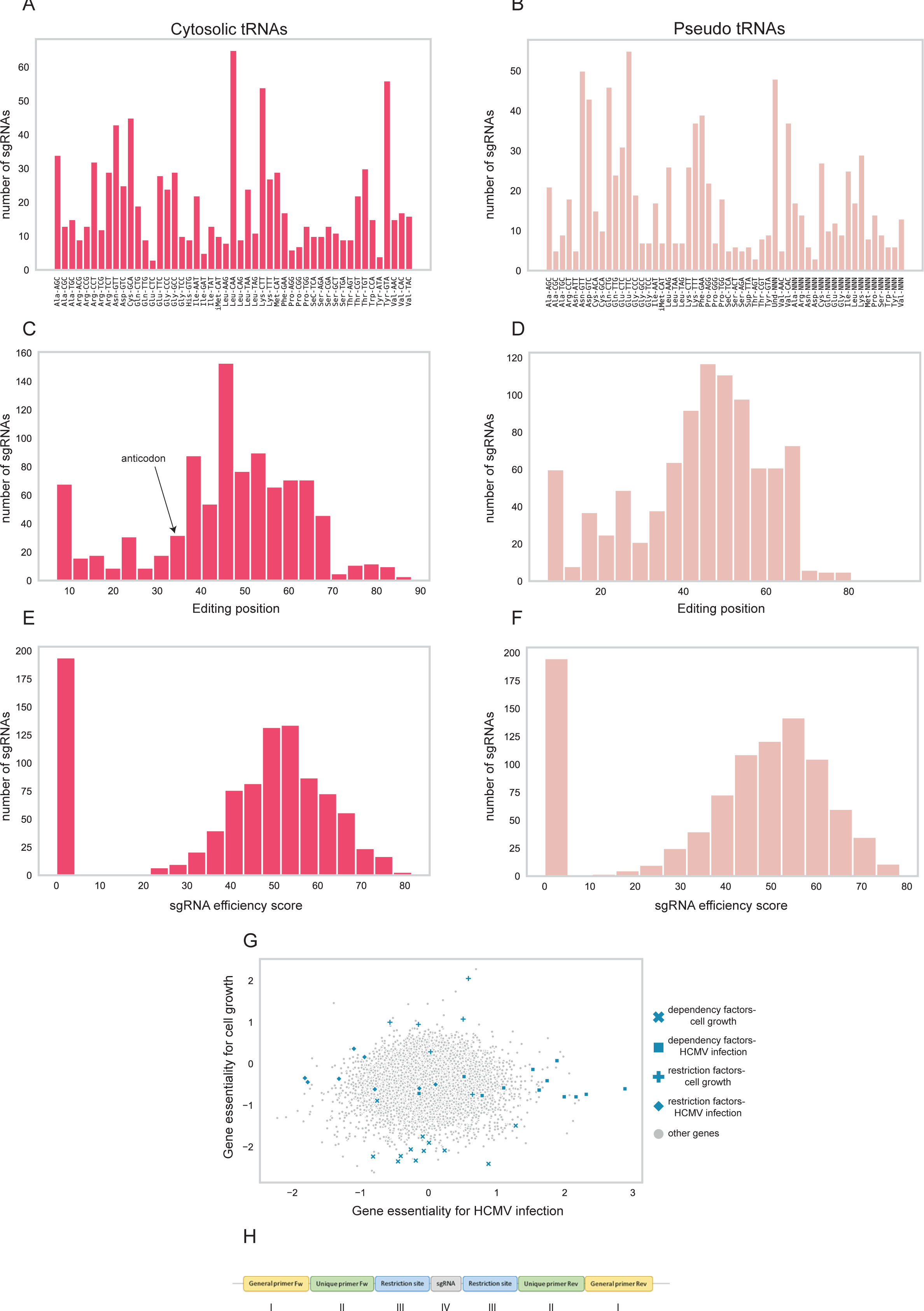
A-B| Number of sgRNAs (y-axis) targeting each tRNA family (x-axis) of A|functional cytosolic tRNA or a B| pseudo-tRNA. C-D| Histograms describing the editing positions of sgRNAs that target C| functional cytosolic tRNA families D| pseudo-tRNA families. The arrow points to the location of the anticodon in the tRNA gene. E-F| Histograms describing the sgRNA efficiency score (calculated by the sgRNA design tool of Benchling ([Biology Software]. (2022). Retrieved from https://benchling.com)) of sgRNAs targeting E| functional cytosolic tRNA families F| pseudo-tRNA families. G| Gene essentiality from a published CRISPR screen of HFFs infected with HCMV (Hein & Weissman, 2022). The colored genes are the ones that were chosen to serve as control genes in the tRNA-CRISPR library, and their marker shape correspond to the marker describes each sub-library in Figure 5B legend. H| Description of The final oligo design contains: I-General primers (common to all sgRNA variants), II-sub-library specific primers (to allow selective amplification of each sub-library), III-restriction sites for BsmB1 restriction enzyme, IV-unique sgRNA.

**Figure S8.**
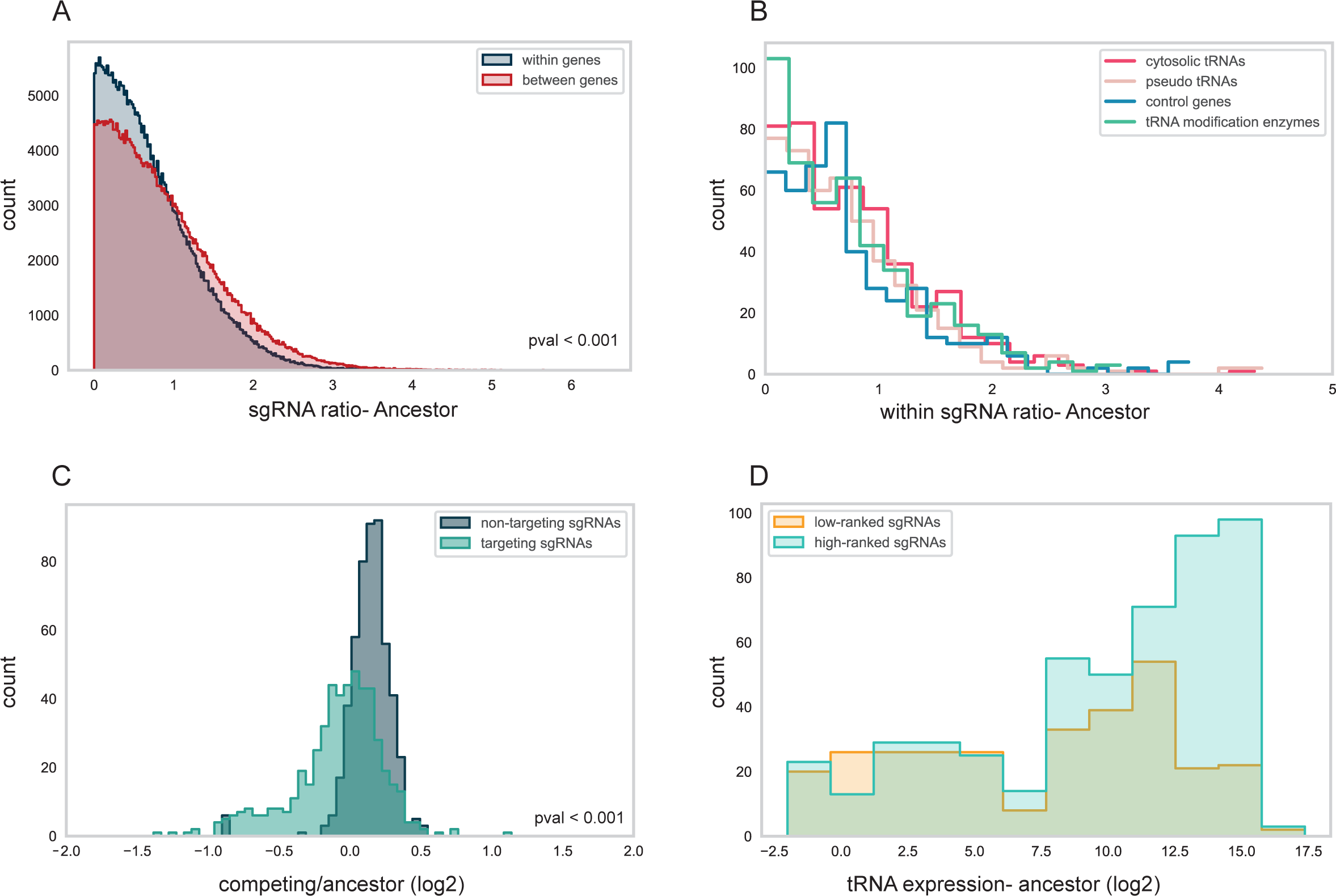
A| Histogram of the ratios between normalized read counts of sgRNAs that target the same gene (“within genes”, blue) and sgRNAs that target different genes (“between genes”, red). Data is shown for the ancestor population. Wilcson rank-sum test p-value < 0.001 B| Histograms of the ratios between normalized read counts of sgRNAs that target the same gene (“within genes”) for different sub-libraries in the ancestor population. C| Histograms comparing sgRNA enrichment in competing relative to ancestor cells between targeting sgRNAs (light green) and non-targeting sgRNAs (dark green). Wilcson rank-sum test p-value < 0.001 D| Histograms describing the tRNA expression (log2) of the targeted tRNAs by highly ranked sgRNAs (turquoise) and lowly ranked sgRNAs (orange). Wilcson rank-sum test p-value < 0.001.

